# Brain Regions Involved in Object-Location Memory Across the Human Lifespan: A Systematic Review and Activation Likelihood Estimation Meta-Analysis of Task-Based fMRI

**DOI:** 10.64898/2026.07.01.735849

**Authors:** Anna Elisabeth Fromm, Mohamed Abdelmotaleb, Freya Olschewski, Jakub Limanowski, Marcus Meinzer, Agnes Flöel, Daria Antonenko

**Author notes:** Corresponding authors, E-Mail addresses and, Department of Neurology, Universitätsmedizin Greifswald, Ferdinand-Sauerbruch-Straße, 17475 Greifswald, Germany. Contributed equally (Anna Elisabeth Fromm, Mohamed Abdelmotaleb, Agnes Flöel, Daria Antonenko).

## Abstract

**Background:** The ability to remember object locations in real life is a fundamental cognitive process that supports goal-directed behavior and is particularly vulnerable to aging and neurodegenerative disease. Despite a growing body of functional magnetic resonance imaging (fMRI) research on object-location memory (OLM), the neural substrates of establishing and retrieving location information are largely unknown.

**Objective:** This systematic review and coordinate-based meta-analysis aimed to identify brain regions consistently activated during OLM in healthy adults, primarily for encoding and – on an exploratory basis – for retrieval, and to characterize age-related differences in OLM-related neural activity.

**Methods:** A systematic search was conducted across three databases (PubMed, PsycInfo, Cochrane Library) up to February 2026. Studies employing task-based fMRI during the encoding and retrieval of object-location associations in healthy adults were eligible. Age-related differences in OLM-related brain activity were examined via narrative synthesis. An activation likelihood estimation (ALE) meta-analysis was performed on studies reporting stereotactic peak coordinates. The review was pre-registered on PROSPERO (CRD420251023695).

**Results:** Twenty-one studies comprising 637 participants were included in the systematic review, with 12 studies being eligible for the encoding ALE meta-analysis. The retrieval ALE meta-analysis was not possible due to the limited number of included studies and reported foci. The systematic review indicated that OLM encoding consistently recruited bilateral fusiform gyri and parahippocampal cortices, with additional engagement of parietal and prefrontal regions across individual studies, whereas OLM retrieval recruited mainly the hippocampus and precuneus. The coordinate-based ALE meta-analysis revealed two significant clusters of activation during OLM encoding: a left-lateralized cluster encompassing the fusiform gyrus, parahippocampal gyrus, and inferior temporal gyrus (peak MNI: −28, −38, −16), and a right-hemisphere cluster spanning the parahippocampal gyrus and fusiform gyrus (peak MNI: 30, −46, −16). Age-related differences, based on a small number of studies with direct age comparison, pointed toward reduced activity in posterior cortical regions coupled with increased activity in prefrontal and midline regions. Additionally, younger adults showed greater hippocampal activation for successful than unsuccessful spatial retrieval, whereas older adults showed the opposite pattern.

**Conclusion:** The systematic review and meta-analysis identify the fusiform gyri and parahippocampal cortices as the most reliably activated regions during OLM encoding, locating OLM formation primarily within the ventral visual-to-medial-temporal processing stream. Retrieval additionally engaged the hippocampus and precuneus, consistent with their established roles in episodic memory. Age-related differences included reduced posterior cortical encoding activity in older adults, a reversal of the hippocampal activation pattern during retrieval, and weaker suppression of midline regions during task performance. The identified encoding pathway may inform targeted network-level interventions such as non-invasive brain stimulation to counteract cognitive decline in aging and neurodegenerative disease.

**Highlights:**

- First systematic review and coordinate-based meta-analysis of task-based fMRI during encoding and retrieval of object-location memories (OLMs).
- Activation likelihood estimation (ALE) identified bilateral fusiform gyrus and parahippocampus as the most consistent encoding regions, supporting a ventral-dominant input pathway.
- Hippocampus and precuneus were the most consistent OLM retrieval regions, suggesting an encoding-retrieval dissociation.
- Age-related encoding differences centered on reduced fusiform and parahippocampal subsequent-memory effects, with an altered anterior-hippocampal response in older adults.

## Introduction

Spatial memory spans a range of abilities, from navigation and map-like representations of large-scale environments to memory for the positions of individual objects. Within this domain, object-location memory (OLM) refers specifically to remembering where objects are located, a form of episodic memory that depends on binding object identity to spatial context and is distinct from large-scale navigation (Postma et al., 2004, 2008). This memory is central to goal-directed behavior throughout life, as we must continually encode and update the changing locations of objects in everyday settings, such as recalling where a car was parked or where a key or wallet was placed (Postma et al., 2008; Zimmermann & Eschen, 2017).

Several neurocognitive frameworks converge on a set of candidate neural systems supporting object-location memory. According to the neurocognitive frameworks proposed by Postma and colleagues (2004, 2008), OLM may be decomposed into three cognitive subprocesses: object processing (the identification of items), spatial-location processing (the encoding of their positions), and object-location binding (the integration of objects with their respective locations). During encoding, object identity and spatial position must be simultaneously maintained until consolidated, i.e., objects being remembered within their initial spatial and temporal context (Gillis et al., 2016; Postma et al., 2008). Postma and colleagues mapped these subprocesses onto partly distinct neural systems: ventral occipitotemporal regions supporting object identification, dorsal parieto-occipital regions supporting spatial localization, and medial temporal lobe (MTL) structures supporting their integration (Postma et al., 2004, 2008).

Subsequent models of episodic memory have refined this mapping. Episodic memory models (Diana et al., 2007; Ranganath & Ritchey, 2012) attribute object processing to the perirhinal cortex, contextual and spatial processing to the parahippocampal cortex, and relational binding to the hippocampus, locating these structures within a broader posterior-medial network, including retrosplenial cortex, posterior cingulate, and precuneus, that is functionally differentiated along the MTL structures. More recent connectivity-based accounts (Rolls, 2024; Rolls et al., 2024) further propose that object identity (“what”) reaches the hippocampus through a ventrolateral occipitotemporal pathway and viewed spatial location (“where”) information primarily via ventromedial occipitotemporal pathways, with the dorsal parietal stream contributing self-motion update rather than primary spatial input. Together, these frameworks converge on a distributed encoding network spanning the ventral and dorsal visual streams and the MTL, while differing in the specific pathways and structures they emphasize. Critically, none of these models has yet been tested against the quantitative convergence of the imaging literature, and all are framed primarily around encoding. Although they partly addressed retrieval – assigning it variously to hippocampal recollection (Postma et al., 2008), hippocampal pattern completion (Rolls, 2024), or posterior-medial contextual reconstruction (Ranganath & Ritchey, 2012) – none specifies whether OLM encoding and retrieval recruit overlapping or dissociable networks, a distinction that remains largely untested for OLM.

OLM is particularly sensitive to the effects of aging. Among many cognitive abilities that decline in healthy aging, including working memory, processing speed, and executive function (Lövdén et al., 2020; Nyberg et al., 2012), OLM is particularly vulnerable because it depends on the coordinated engagement of these multiple subsystems (Gillis et al., 2016; Postma et al., 2008). Behaviorally, older adults show consistent and pronounced deficits in precise spatial localization and object-location binding, whereas object recognition itself is relatively spared (Castegnaro et al., 2022; Meulenbroek et al., 2010; Sapkota et al., 2020). This dissociation has been interpreted as a selective impairment of MTL binding mechanisms rather than a generalized memory deficit (Hampstead et al., 2011; Sapkota et al., 2020; Snytte et al., 2024). Moreover, OLM deficits extend to mild cognitive impairment (MCI) and Alzheimer’s disease (AD), underscoring the translational value of clarifying its neural basis (Castegnaro et al., 2022; Zokaei et al., 2020; Hampstead et al., 2018; Külzow et al., 2014; Kessels et al., 2010). Despite this clinical and ecological relevance, OLM has received less attention than other episodic memory domains, partly because of the difficulty of measuring and operationalizing it (Zimmermann & Eschen, 2017). The literature lacks a standardized assessment framework, and experimental paradigms differ substantially in task design, stimulus materials, retention intervals, and the cognitive operations they recruit (Cona & Scarpazza, 2019; Van Asselen et al., 2008; Zimmermann & Eschen, 2017). This heterogeneity is not merely methodological, it also reflects the multidimensional nature of OLM, as different paradigms probe partially dissociable cognitive and neural mechanisms. Key dimensions include the retention interval (working vs. long-term OLM), the memory stage targeted (encoding vs. retrieval), the spatial reference frame (allocentric vs. egocentric), and the encoding mode (implicit vs. explicit) (Gillis et al., 2016; Postma et al., 2008; Zimmermann & Eschen, 2017).

This heterogeneity is also reflected in fMRI research, where OLM has been operationalized using paradigms spanning spatial memory (Cona & Scarpazza, 2019), object-location memory (Postma et al., 2004; Zimmermann & Eschen, 2017), and source/contextual memory (Davachi, 2006; Kim & Voss, 2019; Slotnick et al., 2003). Existing syntheses reflect these divergent framings. Cona and Scarpazza (2019) conducted a meta-analysis of spatial cognition that pooled long-term spatial memory and navigation studies into a single category, thereby combining paradigms (mental navigation in virtual environments, scene memory, OLM) that may engage partially distinct circuits. By contrast, Zimmermann and Eschen (2017) focused specifically on episodic OLM in young adults and reviewed studies that contrasted OLM with object-memory or location-memory conditions across lesion, fMRI, and PET modalities. However, because their synthesis was qualitative and restricted to studies using this specific subtractive design, it offered a more selective account of the neural mechanisms underlying OLM. To date, no coordinate-based meta-analysis has isolated the neural correlates of OLM specifically.

## The present review

Although many functional imaging studies have investigated OLM, the brain networks supporting it remain incompletely characterized. To our knowledge, this is the first systematic review and coordinate-based meta-analysis of the neural correlates of OLM in healthy adults using task-based fMRI. We synthesized published task-based fMRI evidence on the formation of object-location associations, complemented by an analysis of retrieval-related activity, with two objectives: first, to characterize patterns of task-dependent fMRI activation during OLM encoding and retrieval in healthy younger (18–45 years) and older (55–80 years) adults through a systematic review of all eligible studies; and second, to identify brain regions showing consistent activation across studies using the activation likelihood estimation (ALE) approach (Eickhoff et al., 2009; Turkeltaub et al., 2012). Coordinate-based meta-analysis and ALE in particular treats reported peaks as three-dimensional Gaussian probability distributions and tests, voxel-wise, whether convergence across studies exceeds chance. ALE is well suited to the OLM literature because (a) studies differ widely in stimuli, contrasts, and analysis pipelines but report peaks in common standard spaces; and (b) the construct is sufficiently broad that no single paradigm dominates.

To pursue these aims, we applied a two-tier inclusion approach. All studies meeting the paradigmatic and population criteria outlined below were included in the systematic review (Tier 1), regardless of analysis method or whether encoding or retrieval was performed inside the scanner. From this broader set, studies additionally reporting whole-brain General Linear Model (GLM) contrasts with in-scanner encoding and peak coordinates in standard space were included in the main ALE meta-analysis (Tier 2). In addition, a complementary ALE meta-analysis of OLM retrieval was intended, but too few studies reported usable whole-brain retrieval coordinates to meet the minimum required for a coordinate-based analysis; retrieval was therefore synthesized qualitatively. This design allowed us to maximize the extent of synthesized evidence while preserving the methodological precision required for coordinate-based neuroimaging meta-analysis.

Together, these analyses aim to advance understanding of the neural mechanisms supporting OLM encoding and retrieval across early and late adulthood, with the potential to provide an empirical basis for spatially targeted interventions, in particular non-invasive brain stimulation (NIBS), which can act on the specific network nodes that a coordinate-based meta-analysis identifies, in healthy aging and neurodegenerative disease.

## Methods

### Search strategy and study selection

This systematic review and meta-analysis was pre-registered on PROSPERO (CRD420251023695; registered July 2025). It was conducted in accordance with the PRISMA 2020 guidelines (Page et al., 2021), and the recommendations for neuroimaging meta-analyses (Müller et al., 2018). The literature search was conducted between July 2025 and February 2026. A systematic literature search was conducted in three electronic databases: PubMed, PsycInfo, and Cochrane Library. In all databases, the following search string was applied: (object OR item) AND (location OR position OR spatial) AND (memory OR learning OR associat* OR binding) AND (fMRI OR “functional magnetic resonance imaging” OR “functional neuroimaging” OR “functional MRI”) AND (task OR “task-dependent” OR “task-related” OR “experimental task” OR paradigm). In PubMed, additional filters were applied to restrict the results to (a) human studies, (b) adults aged 19 years and older, and (c) publications written in English. In PsycInfo, the search was further limited to (a) human participants, (b) age groups including young adulthood (18-29 years), thirties (30-39 years), middle age (40-64 years) and aged (65 years and older), (c) peer-reviewed journal articles, (d) studies excluding animal studies, and (e) empirical studies, with literature reviews, interviews, and mathematical models excluded. No additional filters were applied in the Cochrane Library. Additional records were identified through forward and backward citation screening of included articles.

Screening was performed in two sequential phases by three independent researchers (A.E.F., M.A., F.O.). In the first phase, F.O. and A.E.F. independently evaluated the titles and abstracts of all retrieved records to determine potential eligibility. Records deemed potentially relevant were retrieved in full text and, in the second phase, independently assessed against the inclusion criteria by M.A. and F.O. After each phase, reviewers compared their decisions, and any discrepancies were resolved through discussion and consensus among all three reviewers.

Studies were eligible for inclusion in the systematic review if they met all of the following criteria: (a) recruited healthy adult participants (≥ 18 years); (b) employed a task-based fMRI paradigm assessing OLM formation, operationally defined as the active encoding and/or retrieval of associations between objects and their spatial locations; and (c) were published in a peer-reviewed journal and reported novel data on OLM.

Studies were excluded from both the systematic review and the meta-analysis if they (a) exclusively reported patient samples without a separate group of healthy adults; (b) employed paradigms targeting spatial navigation (e.g., trajectory-based navigation in a virtual environment) or working memory rather than discrete, long-term episodic object-location memory; or (c) used PET or other non-fMRI neuroimaging modalities.

From the studies included in the systematic review, an additional set of criteria was applied to identify those eligible for the ALE meta-analysis. Studies were included in the meta-analysis if they employed a whole-brain univariate GLM approach with at least one contrast against a control or baseline condition. Because control conditions ranged from low-level fixation baselines to demanding active control tasks, which differ in the specificity of the resulting contrast, we recorded the control condition for each experiment and considered its potential influence on convergence in the Limitations. Further eligibility criteria for ALE were that studies reported results as standardized stereotactic coordinates (Talairach & Tournoux, 1988) or Montreal Neurological Institute (MNI) space or statistical maps with peak activations, applied threshold, and correction method explicitly stated. Studies were retained in the systematic review but excluded from the ALE analysis if they (a) reported only functional connectivity or multivariate pattern analyses (e.g., partial least squares) of OLM-related networks, or (b) presented only descriptive findings without quantifiable activation data suitable for coordinate-based meta-analysis.

### Data extraction

For each study, we separately extracted brain regions associated with encoding or retrieval activation and brain regions showing age-related differences, including the specific contrasts and statistical methods used. Given the limited number of studies that directly compared younger and older adults during encoding (“cross-age” designs), studies comparing age groups during retrieval were also considered for the systematic review. For a more detailed characterization of the sample, the age (mean, including standard deviation or range), sex ratio (in percent), and MRI parameters (magnetic field strength) were extracted. Additionally, a short description of the task was provided. This information is displayed in Table 2.

For examining age-related differences, studies were grouped into younger (18–45 years) and older (55–80 years) adult samples. These categories were chosen primarily to reflect the age ranges reported in the included studies and to facilitate comparisons across experiments. Nevertheless, the resulting age split is broadly consistent with conventions in cognitive aging and neuroimaging research, where older adulthood is commonly defined from approximately 55–60 years onward (Park & Reuter-Lorenz, 2009; Cabeza et al., 2018).

The final sample comprised 21 studies (systematic review), of which 12 met the criteria for the meta-analysis.

### ALE meta-analysis

Coordinate-based meta-analyses were conducted using the revised ALE algorithm implemented in GingerALE (version 3.0.2; BrainMap) (Eickhoff et al., 2012; Turkeltaub et al., 2012). ALE was performed for experiments investigating OLM encoding of object-location associations. All analyses were carried out in MNI152 space, with the gray-matter search volume defined by the ICBM tissue probability maps implemented in GingerALE. Coordinates originally reported in Talairach space were converted to MNI space using the Lancaster tal2icbm transformation, which reduces the disparity between coordinate systems relative to earlier transforms (Laird et al., 2010; Lancaster et al., 2007). Activation foci were extracted from each included study and entered manually into GingerALE.

For each experiment, ALE models each reported focus as the center of a 3D Gaussian probability distribution representing the spatial uncertainty of the peak location, with the full-width at half maximum (FWHM) derived from empirical estimates of between-subject and between-template variability and scaled inversely with the square root of the experiment’s sample size (Eickhoff et al., 2009). Voxel-wise modeled activation (MA) values were then computed by taking the maximum probability across all foci within each experiment, thereby controlling for within-experiment effects (Turkeltaub et al., 2012). Convergence of activation across experiments was quantified by combining the MA maps to produce voxel-wise ALE scores, with each score reflecting the above-chance convergence of reported peaks across studies (Eickhoff et al., 2009).

Statistical inference followed the recommendations of the large-scale ALE simulation study by Eickhoff et al. (2016) and neuroimaging meta-analysis guidelines (Müller et al., 2018). A cluster-forming threshold of p < 0.001 (uncorrected) was applied at the voxel level, and surviving clusters were tested against a cluster-level family-wise error (FWE) threshold of p < 0.05. The cluster-level null distribution of maximum cluster sizes was estimated by Monte Carlo simulation with 10,000 permutations: for each iteration, foci were randomly relocated within the gray-matter mask while preserving, for each experiment, the original number of foci and the sample-size-dependent FWHM; an ALE analysis was performed on the simulated dataset; and the same cluster-forming threshold (p < 0.001 uncorrected) was applied. Only clusters exceeding the simulation-derived minimum cluster size at pFWE < 0.05 were considered significant. The analysis was carried out in MNI152 space using GingerALE’s more conservative gray-matter mask (228,483 within-brain voxels). Thresholded ALE maps were overlaid on the anatomical MNI template and visualized using MRIcroGL (version 1.2; Rorden, 2025). Anatomical labels were obtained from the Talairach Daemon (Lancaster et al., 2000), as implemented in GingerALE, and cross-referenced against the Harvard-Oxford (Desikan et al., 2006) and Jülich-Brain (Amunts et al., 2020) probabilistic atlases for cerebral peaks and the SUIT atlas (Diedrichsen et al., 2009) for cerebellar peaks.

## Results

### 3.1 Study selection and sample characteristics

A total of 21 studies were included in this systematic review and meta-analysis, with 14 studies exclusively investigating the effects in healthy young adults (Abdelmotaleb et al., 2025; Brodt et al., 2018; Büchel et al., 1999; Cansino, 2002; De Rover et al., 2008; Frings et al., 2006; Gillis et al., 2016; Hales & Brewer, 2013; Kim & Voss, 2019; Rolls et al., 2024; Ross & Slotnick, 2008; Slotnick et al., 2003; Sommer, Rose, Gläscher, et al., 2005; Sommer, Rose, Weiller, et al., 2005), four studies exclusively investigating the effects in healthy older adults (Hampstead et al., 2011, 2016; Rabipour et al., 2020, 2021) and three studies comparing the effects in young vs. older adults (Gould et al., 2003; Kukolja et al., 2009; Meulenbroek et al., 2010). Across these studies, 637 participants were included (N_OA_ = 334, including older adult subsamples from mixed-age studies). Because participants are counted at the level of the publication, this total includes samples re-analyzed across more than one report: Hampstead et al. (2016) re-analyzed the same 16 older adults as Hampstead et al. (2011); the young-adult sample of De Rover et al. (2008) (N=20) overlaps with that of Meulenbroek et al. (2010). None of these reports contributed to the same ALE, so the meta-analysis is unaffected; the descriptive total should be read as a count of analyzed samples rather than of unique individuals.

Twelve studies reported whole-brain GLM activation contrasts with peak coordinates in standard space during encoding and were therefore eligible for the encoding ALE meta-analysis (Abdelmotaleb et al., 2025; Brodt et al., 2018; Cansino, 2002; Frings et al., 2006; Gillis et al., 2016; Gould et al., 2003; Hales & Brewer, 2013; Hampstead et al., 2011; Kukolja et al., 2009; Ross & Slotnick, 2008; Sommer, Rose, Gläscher, et al., 2005; Sommer, Rose, Weiller, et al., 2005). Of these, five scanned and analyzed encoding only (Abdelmotaleb et al., 2025; Gillis et al., 2016; Hales & Brewer, 2013; Sommer, Rose, Gläscher, et al., 2005; Sommer, Rose, Weiller, et al., 2005), whereas the remaining seven additionally acquired a retrieval or recognition phase (Brodt et al., 2018; Cansino, 2002; Frings et al., 2006; Gould et al., 2003; Hampstead et al., 2011; Kukolja et al., 2009; Ross & Slotnick, 2008); in every case, only the encoding contrast was entered into the ALE. The remaining nine studies were not ALE-eligible and contributed to the narrative synthesis only. Three scanned retrieval exclusively (De Rover et al., 2008; Meulenbroek et al., 2010; Slotnick et al., 2003); four applied connectivity-based analyses of OLM networks (Büchel et al., 1999; Hampstead et al., 2016; Kim & Voss, 2019; Rolls et al., 2024); and two used multivariate partial least squares (PLS) modelling instead of univariate GLM (Rabipour et al., 2020, 2021). Study characteristics and fMRI findings of the included studies are presented in Table 1 and Table 2, respectively.

**Table 1:**
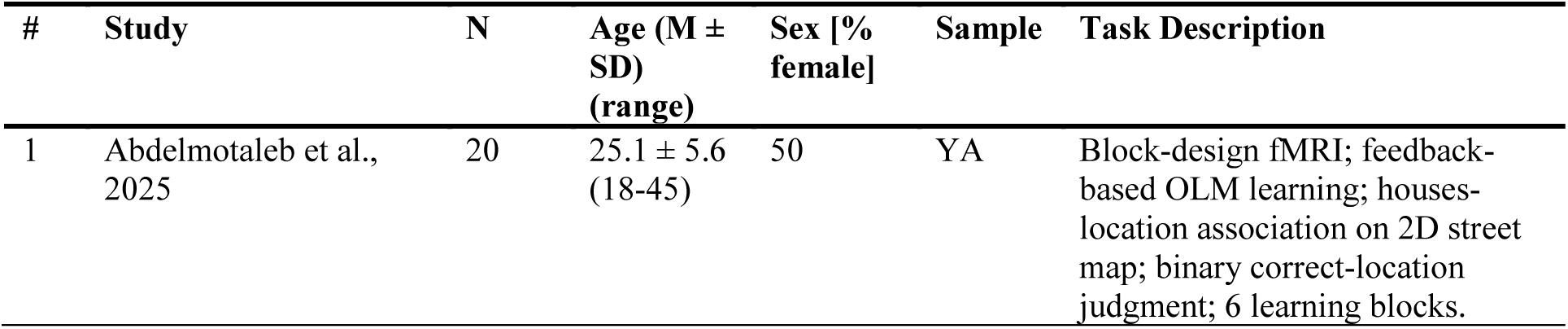
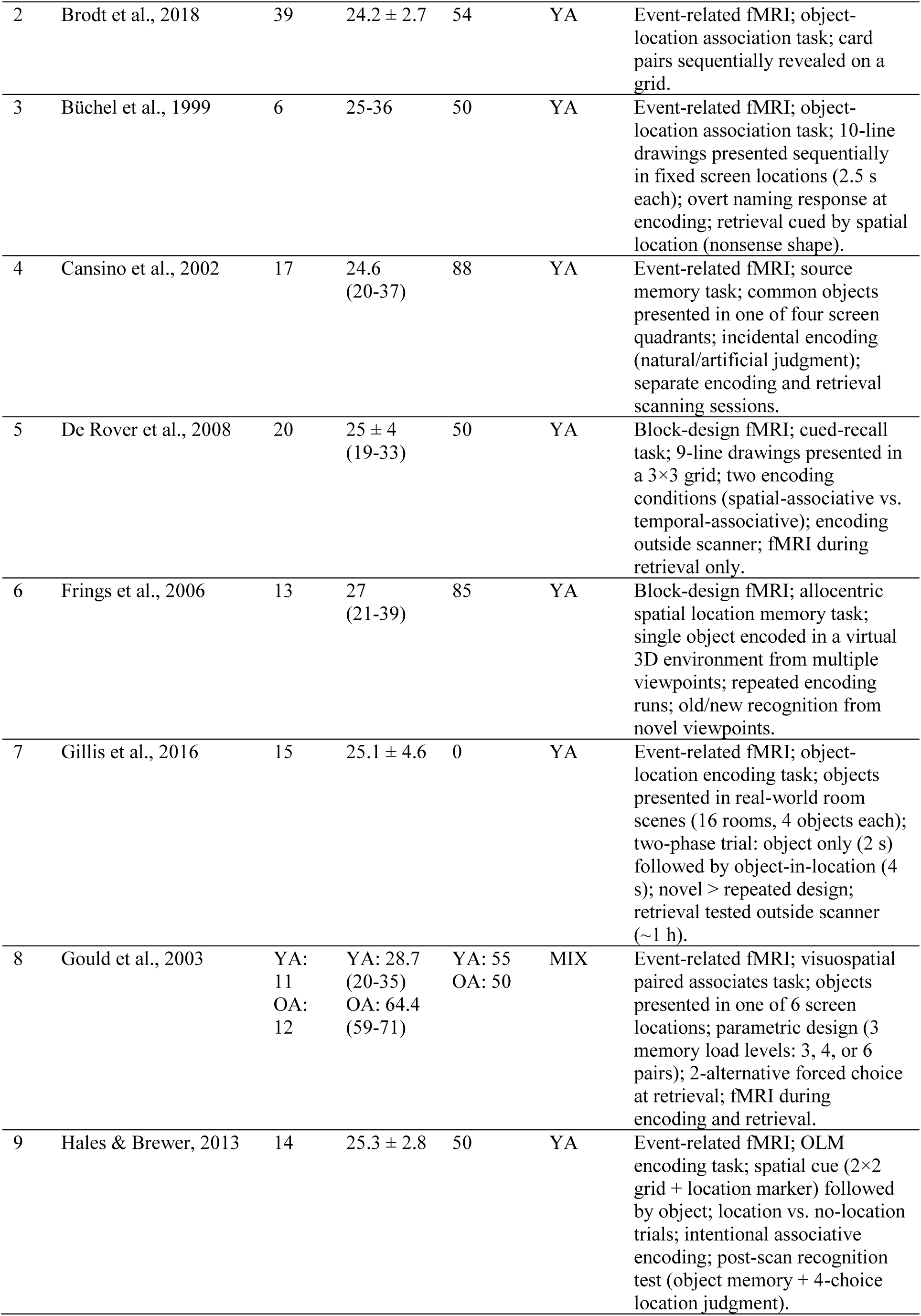
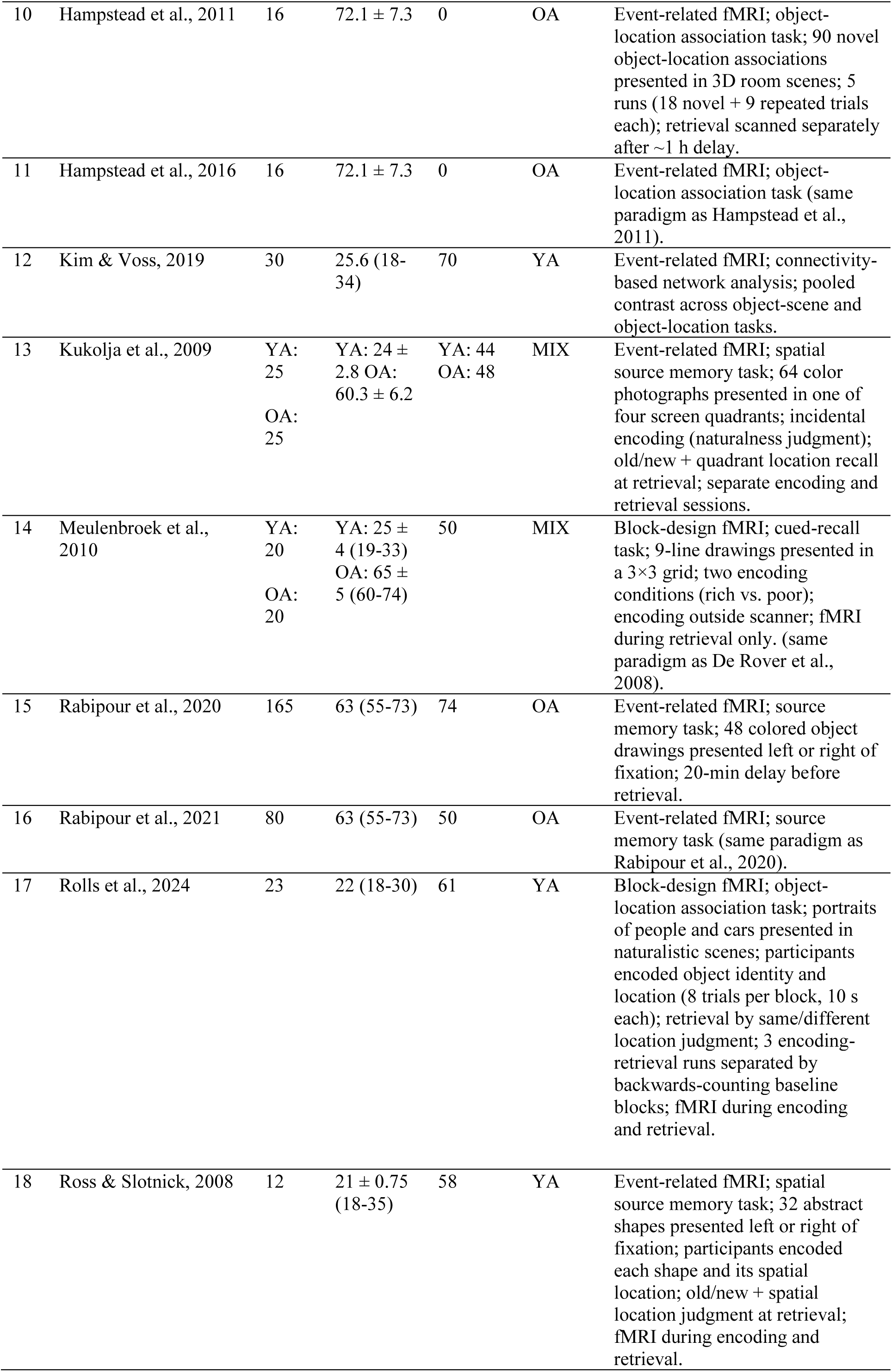
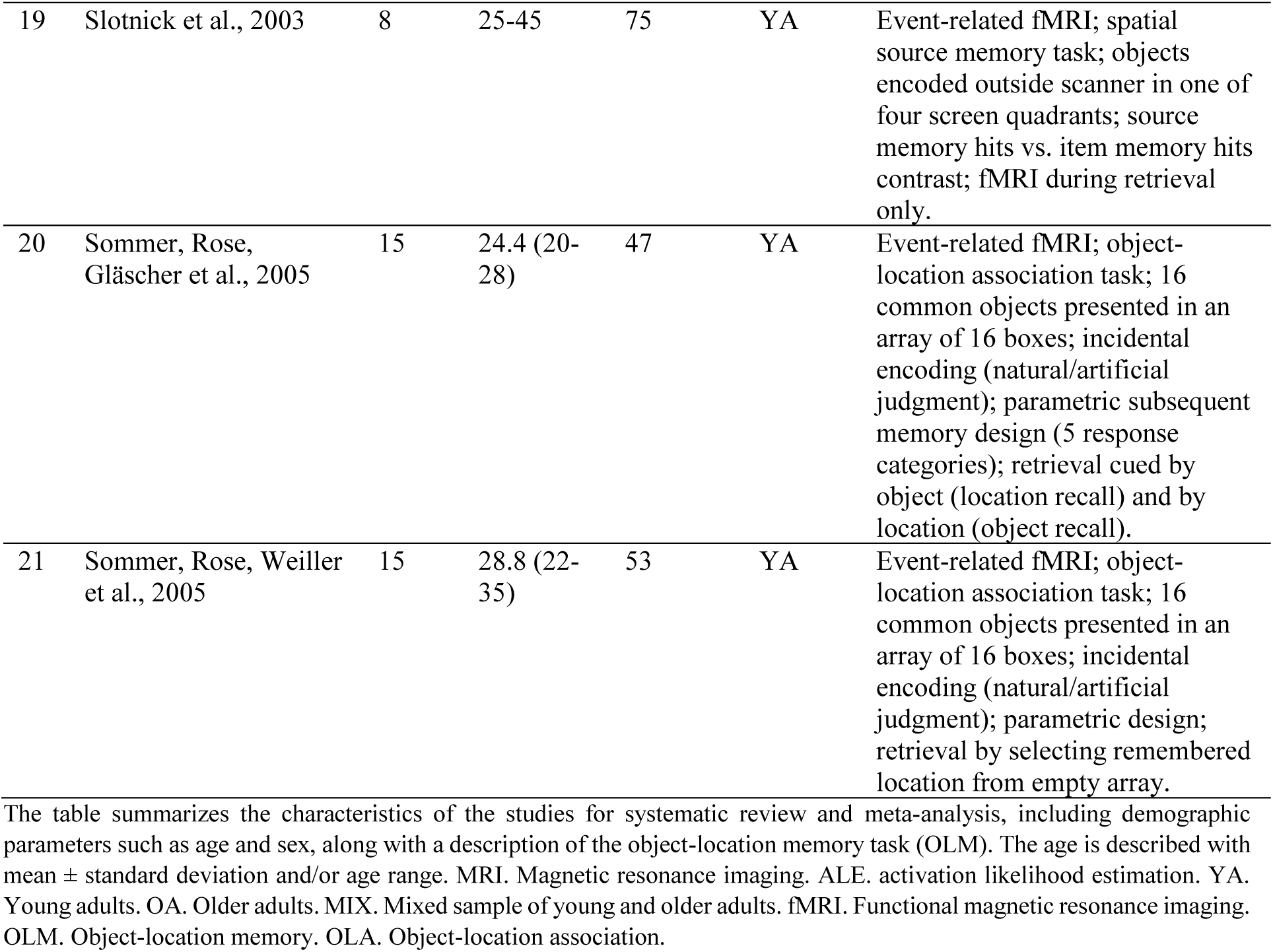
Study characteristics of included studies for the systematic review and meta-analysis.

| # | Study | N | Age (M ± SD)<br>(range) | Sex [% female] | Sample | Task Description |
| --- | --- | --- | --- | --- | --- | --- |
| 1 | Abdelmotaleb et al., 2025 | 20 | 25.1 ± 5.6<br>(18-45) | 50 | YA | Block-design fMRI; feedback-based OLM learning; houses-location association on 2D street map; binary correct-location judgment; 6 learning blocks. |
| 2 | Brodt et al., 2018 | 39 | 24.2 ± 2.7 | 54 | YA | Event-related fMRI; object-location association task; card pairs sequentially revealed on a grid. |
| 3 | Büchel et al., 1999 | 6 | 25-36 | 50 | YA | Event-related fMRI; object-location association task; 10-line drawings presented sequentially in fixed screen locations (2.5 s each); overt naming response at encoding; retrieval cued by spatial location (nonsense shape). |
| 4 | Cansino et al., 2002 | 17 | 24.6<br>(20-37) | 88 | YA | Event-related fMRI; source memory task; common objects presented in one of four screen quadrants; incidental encoding (natural/artificial judgment); separate encoding and retrieval scanning sessions. |
| 5 | De Rover et al., 2008 | 20 | 25 ± 4<br>(19-33) | 50 | YA | Block-design fMRI; cued-recall task; 9-line drawings presented in a 3×3 grid; two encoding conditions (spatial-associative vs. temporal-associative); encoding outside scanner; fMRI during retrieval only. |
| 6 | Frings et al., 2006 | 13 | 27<br>(21-39) | 85 | YA | Block-design fMRI; allocentric spatial location memory task; single object encoded in a virtual 3D environment from multiple viewpoints; repeated encoding runs; old/new recognition from novel viewpoints. |
| 7 | Gillis et al., 2016 | 15 | 25.1 ± 4.6 | 0 | YA | Event-related fMRI; object-location encoding task; objects presented in real-world room scenes (16 rooms, 4 objects each); two-phase trial: object only (2 s) followed by object-in-location (4 s); novel > repeated design; retrieval tested outside scanner (~1 h). |
| 8 | Gould et al., 2003 | YA: 11<br>OA: 12 | YA: 28.7<br>(20-35)<br>OA: 64.4<br>(59-71) | YA: 55<br>OA: 50 | MIX | Event-related fMRI; visuospatial paired associates task; objects presented in one of 6 screen locations; parametric design (3 memory load levels: 3, 4, or 6 pairs); 2-alternative forced choice at retrieval; fMRI during encoding and retrieval. |
| 9 | Hales & Brewer, 2013 | 14 | 25.3 ± 2.8 | 50 | YA | Event-related fMRI; OLM encoding task; spatial cue (2×2 grid + location marker) followed by object; location vs. no-location trials; intentional associative encoding; post-scan recognition test (object memory + 4-choice location judgment). |
| 10 | Hampstead et al., 2011 | 16 | 72.1 ± 7.3 | 0 | OA | Event-related fMRI; object-location association task; 90 novel object-location associations presented in 3D room scenes; 5 runs (18 novel + 9 repeated trials each); retrieval scanned separately after ~1 h delay. |
| 11 | Hampstead et al., 2016 | 16 | 72.1 ± 7.3 | 0 | OA | Event-related fMRI; object-location association task (same paradigm as Hampstead et al., 2011). |
| 12 | Kim & Voss, 2019 | 30 | 25.6 (18-34) | 70 | YA | Event-related fMRI; connectivity-based network analysis; pooled contrast across object-scene and object-location tasks. |
| 13 | Kukolja et al., 2009 | YA: 25<br>OA: 25 | YA: 24 ± 2.8<br>OA: 60.3 ± 6.2 | YA: 44<br>OA: 48 | MIX | Event-related fMRI; spatial source memory task; 64 color photographs presented in one of four screen quadrants; incidental encoding (naturalness judgment); old/new + quadrant location recall at retrieval; separate encoding and retrieval sessions. |
| 14 | Meulenbroek et al., 2010 | YA: 20<br>OA: 20 | YA: 25 ± 4 (19-33)<br>OA: 65 ± 5 (60-74) | 50 | MIX | Block-design fMRI; cued-recall task; 9-line drawings presented in a 3×3 grid; two encoding conditions (rich vs. poor); encoding outside scanner; fMRI during retrieval only. (same paradigm as De Rover et al., 2008). |
| 15 | Rabipour et al., 2020 | 165 | 63 (55-73) | 74 | OA | Event-related fMRI; source memory task; 48 colored object drawings presented left or right of fixation; 20-min delay before retrieval. |
| 16 | Rabipour et al., 2021 | 80 | 63 (55-73) | 50 | OA | Event-related fMRI; source memory task (same paradigm as Rabipour et al., 2020). |
| 17 | Rolls et al., 2024 | 23 | 22 (18-30) | 61 | YA | Block-design fMRI; object-location association task; portraits of people and cars presented in naturalistic scenes; participants encoded object identity and location (8 trials per block, 10 s each); retrieval by same/different location judgment; 3 encoding-retrieval runs separated by backwards-counting baseline blocks; fMRI during encoding and retrieval. |
| 18 | Ross & Slotnick, 2008 | 12 | 21 ± 0.75 (18-35) | 58 | YA | Event-related fMRI; spatial source memory task; 32 abstract shapes presented left or right of fixation; participants encoded each shape and its spatial location; old/new + spatial location judgment at retrieval; fMRI during encoding and retrieval. |
| 19 | Slotnick et al., 2003 | 8 | 25-45 | 75 | YA | Event-related fMRI; spatial source memory task; objects encoded outside scanner in one of four screen quadrants; source memory hits vs. item memory hits contrast; fMRI during retrieval only. |
| 20 | Sommer, Rose, Gläscher et al., 2005 | 15 | 24.4 (20-28) | 47 | YA | Event-related fMRI; object-location association task; 16 common objects presented in an array of 16 boxes; incidental encoding (natural/artificial judgment); parametric subsequent memory design (5 response categories); retrieval cued by object (location recall) and by location (object recall). |
| 21 | Sommer, Rose, Weiller et al., 2005 | 15 | 28.8 (22-35) | 53 | YA | Event-related fMRI; object-location association task; 16 common objects presented in an array of 16 boxes; incidental encoding (natural/artificial judgment); parametric design; retrieval by selecting remembered location from empty array. |

The table summarizes the characteristics of the studies for systematic review and meta-analysis, including demographic parameters such as age and sex, along with a description of the object-location memory task (OLM). The age is described with mean ± standard deviation and/or age range. MRI. Magnetic resonance imaging. ALE. activation likelihood estimation. YA. Young adults. OA. Older adults. MIX. Mixed sample of young and older adults. fMRI. Functional magnetic resonance imaging. OLM. Object-location memory. OLA. Object-location association.

**Table 2:** fMRI findings of the studies included in the systematic review and ALE meta-analysis.

| # | Study | MRI | fMRI analysis (type, correction, contrast, coordinate space) | ALE-eligible | Key reported regions encoding/retrieval |
| --- | --- | --- | --- | --- | --- |
| 1 | Abdelmotaleb et al., 2025 | 3 T | Whole-brain GLM; voxel $p < 0.001$ with FWE cluster correction ($k > 125$); Learning > Control; MNI space. | Yes | **Encoding (Learning > Control):** bilateral temporo-occipital & posterior fusiform gyrus; R hippocampus; bilateral LOC; precuneus; R caudate/putamen; bilateral orbitofrontal cortex; bilateral precentral gyrus; L insula; L ITG. |
| 2 | Brodt et al., 2018 | 3 T | Whole-brain GLM, full-volume FWE peak $p < 0.05$; subsequent memory (enc-correct > enc-incorrect) + experience-dependent increase over runs + performance correlation; precuneus/hippocampus ROI; MNI space. | Yes | **Encoding (subsequent memory, correct > incorrect):** R hippocampus; L STG; L IFG<br>**Retrieval (recall increase over runs / performance correlation):** bilateral precuneus, L posterior cingulate, cerebellum; precuneus activity rose across recall repetitions and persisted > 12 h. |
| 3 | Büchel et al., 1999 | 2 T | Repetition-suppression (time × condition) during OLM encoding + ROI path analysis / SEM of effective connectivity (dorsal vs ventral stream); standard template (MNI-type). | No, Connectivity | **Encoding (associative learning):** repetition-suppression, R: V1, dorsal extrastriate, posterior parietal cortex (PP), posterior inferotemporal/fusiform (ITp).<br>Connectivity results: learning-related increase in effective connectivity from PP (dorsal stream) → ITp/fusiform (ventral stream). |
| 4 | Cansino et al., 2002 | 2 T | Whole-brain GLM, random-effects; voxel $p < 0.001$ uncorrected, $k \geq 5$; Encoding: subsequent correct > incorrect source; Retrieval: correct > incorrect source; Talairach space. | Yes | **Encoding (subsequent source memory):** L SFG, L IFG, bilateral precentral gyrus, paracentral lobule, L IPL/lateral parietal, L superior temporal sulcus, R LOC, bilateral fusiform gyrus, L nucleus accumbens, L cerebellum.<br>**Retrieval (correct > incorrect source):** L medial frontal cortex, R posterior insula, R IPL/lateral parietal, L parahippocampal cortex, R hippocampal formation, R amygdala. |
| 5 | De Rover et al., 2008 | 1.5 T | Whole-brain GLM, random-effects; cluster FWE $p < 0.05$; recall > fixation baseline; Retrieval only scanned, encoding outside scanner; MNI/Talairach space. | No, Retrieval only | **Retrieval (recall > baseline; cortex by BA only):** occipital (BA 17/18/19), parietal (BA 7, 23/31, 39/40), temporal (BA 37), parahippocampal/MTL (BA 36), prefrontal (BA 6/8/9/44/45/46), ACC (BA 6/32), bilateral thalamus & basal ganglia. |
| 6 | Frings et al., 2006 | 1.5 T | Whole-brain GLM; voxel FWE $p < 0.05$; ENC1 > VIEW / ENC1 > COMP1 (perceptual/motor control); encoding and recognition conjunction; MNI space. | Yes | **Encoding (ENC1 > VIEW / COMP1):** bilateral precuneus (BA7; dominant under FWE); also, bilateral cingulate (BA32), L SPL, bilateral IPL, R thalamus, L ITG, R insula, R SFG.<br>**Retrieval / conjunction:** precuneus. |
| 7 | Gillis et al., 2016 | 3 T | Random-effects GLM; voxel $p < 0.05$ + Monte-Carlo cluster correction (189 voxels); Object+Location(novel) > Object+Location(repeated); encoding only (retrieval out of scanner); Talairach space. | Yes | **Encoding (novel > repeated):** bilateral hippocampus; bilateral PHG/entorhinal cortex; bilateral fusiform; bilateral IPS/SPL; precuneus; bilateral MFG/IFG/FEF; ACC; bilateral LOC; retrosplenial cortex; thalamus; striatum (caudate/putamen). |
| 8 | Gould et al., 2003 | 3 T | Whole-brain GLM; FDR (zero-order encoding $q < 0.0001$; Encoding > Retrieval $q < 0.0005$); parametric load; Talairach space. | Yes | **Encoding (zero-order, load-independent, enc > rest):** R precuneus; L MFG; anterior cingulate; bilateral occipital (IOG/MOG/SOG); thalamus; L IFG; bilateral cerebellum.<br>**Encoding > Retrieval:** L IFG, L MFG, R hippocampus, R parahippocampal gyrus. |
| 9 | Hales & Brewer, 2013 | 3 T | Whole-brain GLM; AlphaSim $p < 0.05$, $k \geq 62$; cue-period contrasts (Location > No-location; Correct > Incorrect location); encoding only; Talairach space. | Yes | **Encoding: attempted binding (Location > No-location):** bilateral SPL, R SFG, L MOG.<br>**Encoding: successful binding (correct > incorrect, subsequent memory):** R MFG (BA10), bilateral cingulate (BA32), L cerebellum. |
| 10 | Hampstead et al., 2011 | 3 T | Whole-brain random-effects GLM; within-group FDR $q < 0.05$; successfully encoded Novel > Repeated (healthy older controls); encoding (retrieval scanned separately); Talairach space. | Yes | **Encoding (novel > repeated, healthy older controls):** bilateral hippocampus (anterior/body/posterior); bilateral PHG/entorhinal & perirhinal cortex; bilateral fusiform; bilateral SPL/IPS; precuneus; retrosplenial cortex; posterior cingulate; bilateral MFG/IFG; bilateral thalamus; putamen; amygdala; bilateral MOG; cerebellum. |
| 11 | Hampstead et al., 2016 | 3 T | Effective connectivity (multivariate Granger causality), correct trials; ROIs from novel-correct > repeated conjunction (encoding&retrieval); activation FDR $q \leq 0.05$, paths FDR $p < 0.05$; Talairach space. | No, Connectivity | **Encoding connectivity (healthy older controls):** L inferior frontal junction (IFJ) = primary intrahemispheric driver → projects to L IFJ, vmPFC, posterior cingulate, MTG; L IFJ & L posterior cingulate = co-primary interhemispheric drivers.<br>**Retrieval connectivity (healthy older controls):** R anterior hippocampus = primary driver of intra- & interhemispheric retrieval connectivity (recollection hub); R anterior thalamus & R inferior frontal sulcus = secondary drivers. |
| 12 | Kim & Voss, 2019 | 3 T | Large-scale functional connectivity / graph-theoretic module-allegiance analysis; univariate map ($p < 0.05$ uncorrected) used only for ROI definition; connectivity Bonferroni $P_{corr} < 0.05$; encoding (memory formation); Talairach space. | No, Connectivity | **Encoding (no activation peaks reported):** less within-module coupling of a fronto-parietal module (POS2–POS2) predicted better object-location/source memory; fronto-parietal × default-lymbic coupling (POS2–NEG3) was negatively related to source memory (did not survive Bonferroni). |
| 13 | Kukolja et al., 2009 | 1.5 T | Whole-brain random-effects, age × source judgement; voxel $p < 0.001$ uncorrected, $k \geq 5$; CorSCE $>$ FalsCE (encoding) and CorSCR $>$ FalSCR (retrieval); MNI space. | Yes | **Encoding (CorSCE $>$ FalsCE, subsequent source memory):** L fusiform gyrus (subsequent-memory effect in young, absent in older); L parahippocampal gyrus (encoding activity predicting later correct retrieval; both groups, stronger in older).<br>**Retrieval (CorSCR $>$ FalSCR):** L hippocampal formation, L precuneus, bilateral fusiform gyrus, L SPL, L ACC, L insula, R lingual gyrus, cerebellum.<br>**Age effects:** anterior hippocampal retrieval response reverses in older adults (higher for failure than success). |
| 14 | Meulenbroek et al., 2010 | 1.5 T | Whole-brain random-effects; $2 \times 2$ Age × Condition ANCOVA, performance covariate; cluster-level FWE $p < 0.05$ (voxel-forming $T = 3.20$, $p = 0.001$ uncorrected); cued-recall (retrieval) phase modeled, encoding not analyzed; MNI space. | No, retrieval only study | **Retrieval: main effect of age (Elderly $>$ Young):** L STG (BA21/22/42), R putamen, R insula, R caudate.<br>**Retrieval: interaction (rich $>$ poor environment AND Elderly $>$ Young):** L MTG, L fusiform gyrus, L ITG, R parahippocampal gyrus, R hippocampus, bilateral thalamus (mediodorsal/pulvinar), bilateral globus pallidus, R brainstem.<br>**General cued-recall network (both groups) :** dorsal & ventral visual streams extending into MTL, (pre)motor cortex, DLPFC, basal ganglia. |
| 15 | Rabipour et al., 2020 | 3 T | Multivariate spatiotemporal PLS (Task-PLS, Behavior-PLS); bootstrap ratio $\geq \pm 3.0$–$3.5$ ($\approx p < 0.001$); T-PLS LV1 (encoding vs retrieval); older adults (APOE4); Talairach space. | No, Multivariate | **Encoding (T-PLS LV1 negative salience, enc $>$ retrieval):** R lingual gyrus, L medial frontal cortex, L ITG, L pre/postcentral gyrus, L STG, L inferior occipital gyrus.<br>**Retrieval (T-PLS LV1 positive salience, retrieval $>$ enc):** bilateral precuneus, bilateral posterior cingulate, bilateral lingual gyrus, bilateral superior occipital gyrus, L MFG, medial frontal cortex, thalamus, IPL. |
| 16 | Rabipour et al., 2021 | 3 T | Multivariate spatiotemporal PLS (Task-PLS, Behavior-PLS); bootstrap ratio $\geq +3.28$ / $\leq -3.28$ ($\approx p < 0.001$); T-PLS LV1; older women (PREVENT-AD); Talairach space. | No, Multivariate | **Retrieval (T-PLS LV1 positive salience, retrieval $>$ enc & correct rejections):** bilateral precuneus, R superior occipital gyrus, precentral gyrus, bilateral SFG, bilateral insula, L angular gyrus, L STG/MTG, L posterior cingulate, L MFG.<br>**Encoding (T-PLS LV1 negative salience, enc & CR $>$ retrieval):** R fusiform gyrus, precentral gyrus, R MOG, caudate, L postcentral gyrus, L ITG, medial frontal cortex, R cuneus, L anterior cingulate. |
| 17 | Rolls et al., 2024 | 3 T | Surface-based parcellation (HCP-MMP, 360 regions; HCPex subcortical); activation = RMSSD per parcel; a-priori ROI paired t-tests $p < 0.02$; + effective/functional connectivity; no voxel coordinates; both phases (combined ROI). | No, Parcellations | **OLM $>$ word-pair memory (storage & recall combined):** ventromedial visual stream (VMV1–3, VVC) and medial parahippocampal areas (PHA1–3); intraparietal regions (LIPv, MIP, VIP). Right-lateralized in OLM: ventrolateral face/object stream (FFC, PIT, V8), inferior temporal TE2p, and medial parahippocampal PHA1–3. |
| 18 | Ross & Slotnick, 2008 | 3 T | Whole-brain voxel-wise GLM, random-effects + a-priori MTL/parietal ROI; voxel $p < .01$, $k \geq 137$ (Monte-Carlo corrected $p < .01$); enc-hit-hits > enc-hit-misses (encoding) and old-hit-hits > old-hit-misses (retrieval); Talairach space. | Yes | **Encoding (subsequent source memory):** L parahippocampal cortex, bilateral postcentral gyrus, R supramarginal gyrus, bilateral IPS, R SPL, L fusiform gyrus, L MTG, L IOG, L MOG.<br>**Retrieval (old-hit-hits > old-hit-misses):** bilateral hippocampus, L parahippocampal gyrus, IFG, IPS, SPL, fusiform gyrus, putamen, thalamus. |
| 19 | Slotnick, 2003 | 1.5 T | Whole-brain voxel-wise GLM, random-effects; voxel $p = .01$ ($t > 3$) uncorrected, $k \geq 98$ (Monte-Carlo corrected $p = .05$); Source memory > Item memory (retrieval only); Talairach space. | No, Retrieval only | **Retrieval (source > item memory):** bilateral SFG, L MFG, L IFG, R superior temporal sulcus, L STG. |
| 20 | Sommer, Rose, Gläscher et al., 2005 | 3 T | Whole-brain GLM, parametric subsequent memory, small-volume correction within a-priori 10-mm VOIs; object-cue & location-cue contrasts, conjunction, interaction; encoding (retrieval not scanned); MNI space. | Yes | **Encoding: object cue (subsequent location memory):** R calcarine/V1, R dorsal extrastriate, bilateral SPL, L angular gyrus, bilateral lingual gyrus, bilateral fusiform gyrus, bilateral parahippocampal gyrus, bilateral FEF, L anterior inferior PFC, bilateral anterior MTL, superior colliculi.<br>**Conjunction (predicts BOTH location & object memory):** bilateral parahippocampal cortex, L fusiform gyrus, L anterior inferior PFC.<br>**Interaction (object > location cue; location-specific):** R anterior MTL, L parahippocampal cortex. |
| 21 | Sommer, Rose, Weiller et al., 2005 | 1.5 T | Whole-brain GLM, event-related parametric subsequent-memory (Dm); voxel $p < 0.001$ uncorrected, $k \geq 5$; activity ∝ subsequent location-memory accuracy/confidence; encoding (retrieval not scanned); MNI space. | Yes | **Encoding (subsequent location-memory confidence):** R calcarine/V1–V2, R dorsal extrastriate (~V3a), bilateral SPL, L angular gyrus, bilateral fusiform gyrus, bilateral parahippocampal gyrus, bilateral FEF, L rostral precentral sulcus, L anterior & posterior inferior PFC, R dorsolateral PFC, L anterior cingulate. |

Encoding and retrieval are reported separately wherever a study scanned both phases. Hemisphere: L = left, R = right; “bilateral” = both. ACC, anterior cingulate cortex; AG, angular gyrus; CR, correct rejection; Cun, cuneus; dlPFC, dorsolateral prefrontal cortex; EC, entorhinal cortex; FEF, frontal eye fields; FG, fusiform gyrus; HC, hippocampus; IFG, inferior frontal gyrus; IFJ, inferior frontal junction; IOG/MOG/SOG, inferior/middle/superior occipital gyrus; IPFC, inferior prefrontal cortex; IPL, inferior parietal lobule; IPS, intraparietal sulcus; ITG/MTG/STG, inferior/middle/superior temporal gyrus; LG, lingual gyrus; LOC, lateral occipital complex; LV, latent variable; mPFC/vmPFC, medial/ventromedial prefrontal cortex; MFG/SFG, middle/superior frontal gyrus; MTL, medial temporal lobe; PCC, posterior cingulate cortex; PHC/PHG, parahippocampal cortex/gyrus; PLS, partial least squares; PRC, perirhinal cortex; RMSSD, root-mean-square successive difference; ROI, region of interest; RSC, retrosplenial cortex; SME, subsequent-memory effect; SMG, supramarginal gyrus; SPL, superior parietal lobule; STS, superior temporal sulcus; SVC, small-volume correction; V1, primary visual cortex.

### 3.2 Systematic review

The systematic-review findings are presented in three parts: encoding (3.2.1), retrieval (3.2.2), and age-related differences in both encoding and retrieval (3.2.3).

#### 3.2.1. Brain regions associated with OLM encoding

##### Occipitotemporal cortex

The most consistent encoding finding across all studies was bilateral activation of the fusiform gyrus and parahippocampal cortex. This pattern was observed across different paradigm types, including feedback-based learning (Abdelmotaleb et al., 2025), novel object-location associations (Gillis et al., 2016), parametric load designs (Gould et al., 2003), ecologically oriented encoding in healthy older adults (Hampstead et al., 2011), and multivariate analyses of older adult cohorts (Rabipour et al., 2021). Fusiform activation was identified across paradigms as the dominant ventral-stream response, showing good test-retest reliability (Abdelmotaleb et al., 2025), a strong subsequent memory effect driven by perceptual engagement during encoding (Cansino, 2002), and a parametrically graded relationship with confidence in remembering (Sommer, Rose, Gläscher, et al., 2005; Sommer, Rose, Weiller, et al., 2005). Additionally, parahippocampal activation showed a partially dissociable profile with bilateral posterior parahippocampi predicting memory for object and location components, while the anterior parahippocampal cluster (left) and right anterior MTL were specifically predictive of location recall (Sommer, Rose, Gläscher, et al., 2005). Left parahippocampus activity during encoding further correlated with subsequent retrieval success across age groups, with this relationship being stronger in older adults (Kukolja et al., 2009) and was identified in the encoding-retrieval conjunction (Ross & Slotnick, 2008). Dominant parahippocampal activation at encoding was further reported in Gillis et al. (2016), Gould et al. (2003), Hampstead et al. (2011), and Sommer, Rose, Weiller et al. (2005).

The right lateral occipital complex was the largest encoding cluster in Cansino and colleagues (2002), and bilateral occipital cortex showed reliable activation with good test-retest reliability in two separate OLM encoding sessions (Abdelmotaleb et al., 2025). Occipital activation was additionally reported in Gould et al. (2003) and Ross and Slotnick (2008). Additionally, the right dorsal extrastriate cortex and bilateral lingual gyrus were parametrically modulated by location memory confidence (Sommer, Rose, Gläscher, et al., 2005; Sommer, Rose, Weiller, et al., 2005), and reported to be activated in older adults (Rabipour et al., 2020).

##### Frontal and parietal cortex

Frontal and parietal activation was significant across multiple studies. Hales and Brewer (2013) reported a dissociation between bilateral superior parietal cortex, which was active during any location encoding attempt regardless of subsequent memory outcome, and right middle frontal gyrus, which was selectively engaged for successful object-location binding. Frings et al. (2006) identified bilateral precuneus as the dominant encoding activation under FWE correction. Precuneus activation during encoding was additionally reported in Abdelmotaleb et al. (2025), Gillis et al. (2016), and Gould et al. (2003). Bilateral intraparietal sulcus (IPS) was engaged during source memory encoding in Ross and Slotnick (2008), and bilateral IPS and dorsal precuneus activation was also observed in healthy older adults (Hampstead et al., 2011). Superior parietal cortex was parametrically modulated by location memory confidence, and bilateral frontal eye fields showed a parametric response (Sommer, Rose, Gläscher, et al., 2005; Sommer, Rose, Weiller, et al., 2005). Additionally, frontal activation including the bilateral MFG and IFG was reported (Cansino, 2002; Gillis et al., 2016; Gould et al., 2003; Hampstead et al., 2011).

##### Hippocampus

Bilateral hippocampal activation was reported in novel > repeated object-location contrasts (Gillis et al., 2016), and right hippocampal activation was reported during load-independent encoding (Gould et al., 2003). Hampstead et al. (2011) reported the most extensive hippocampal encoding activation in the present review, with bilateral activation across the full anterior-posterior extent under FDR correction in healthy older adults. The right hippocampus was further identified as a subpeak that uniquely increased across learning stages while other regions showed suppression, consistent with increasing hippocampal engagement as learning progresses (Abdelmotaleb et al., 2025). A right anterior MTL cluster was specifically predictive of location recall but not object identity in an interaction analysis, while bilateral posterior parahippocampal cortex emerged in the conjunction for both memory components (Sommer, Rose, Gläscher, et al., 2005). Several studies reported no hippocampal encoding activation, including Kukolja et al. (2009), Cansino (2002), and Sommer, Rose, Weiller et al. (2005). A hippocampal subsequent memory effect was identified in supplementary data only in Brodt et al. (2018), with no hippocampal activation in their primary recall contrasts.

Moreover, in the connectivity studies examining OLM hippocampal-cortical effective connectivity increased with learning (Büchel et al., 1999), large-scale network integration supported item–context binding (Kim & Voss, 2019), and the hippocampus was the principal driver of retrieval connectivity in older adults, a pattern absents in MCI (Hampstead et al., 2016).

##### Cerebellum

Cerebellar activation was reported across multiple individual studies at encoding (Cansino et al., 2002; Gould et al., 2003; Hales & Brewer, 2013; Hampstead et al., 2011; Ross & Slotnick, 2008). However, no study discussed cerebellar involvement as a primary finding, and its interpretation is complicated by atlas-dependent labeling of ventral occipitotemporal peaks (see Section 4.1).

A schematic summary of the OLM encoding is displayed in Figure 2A.

**Figure 1.**
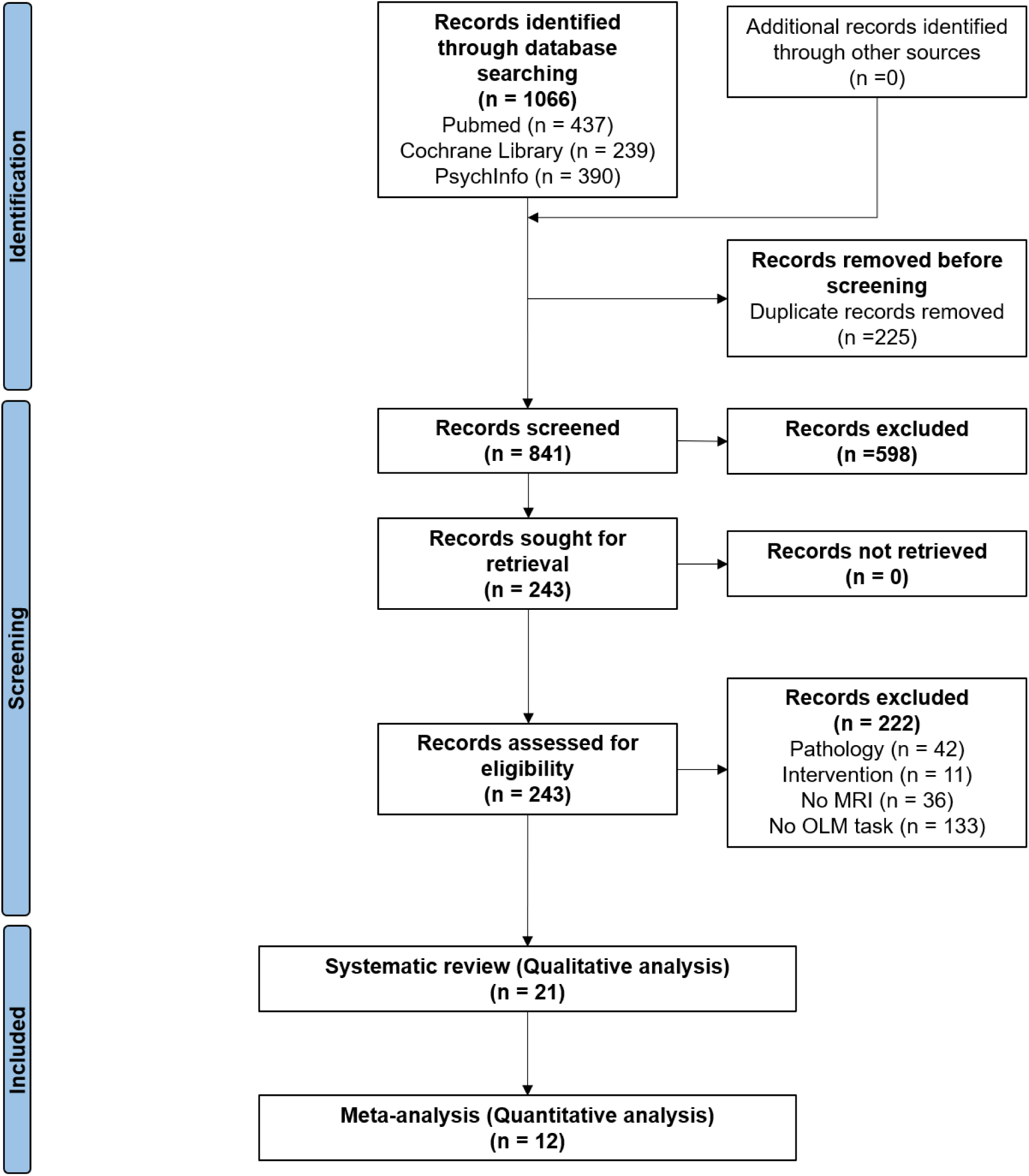
PRISMA flowchart for the study identification procedure.

**Figure 2.**
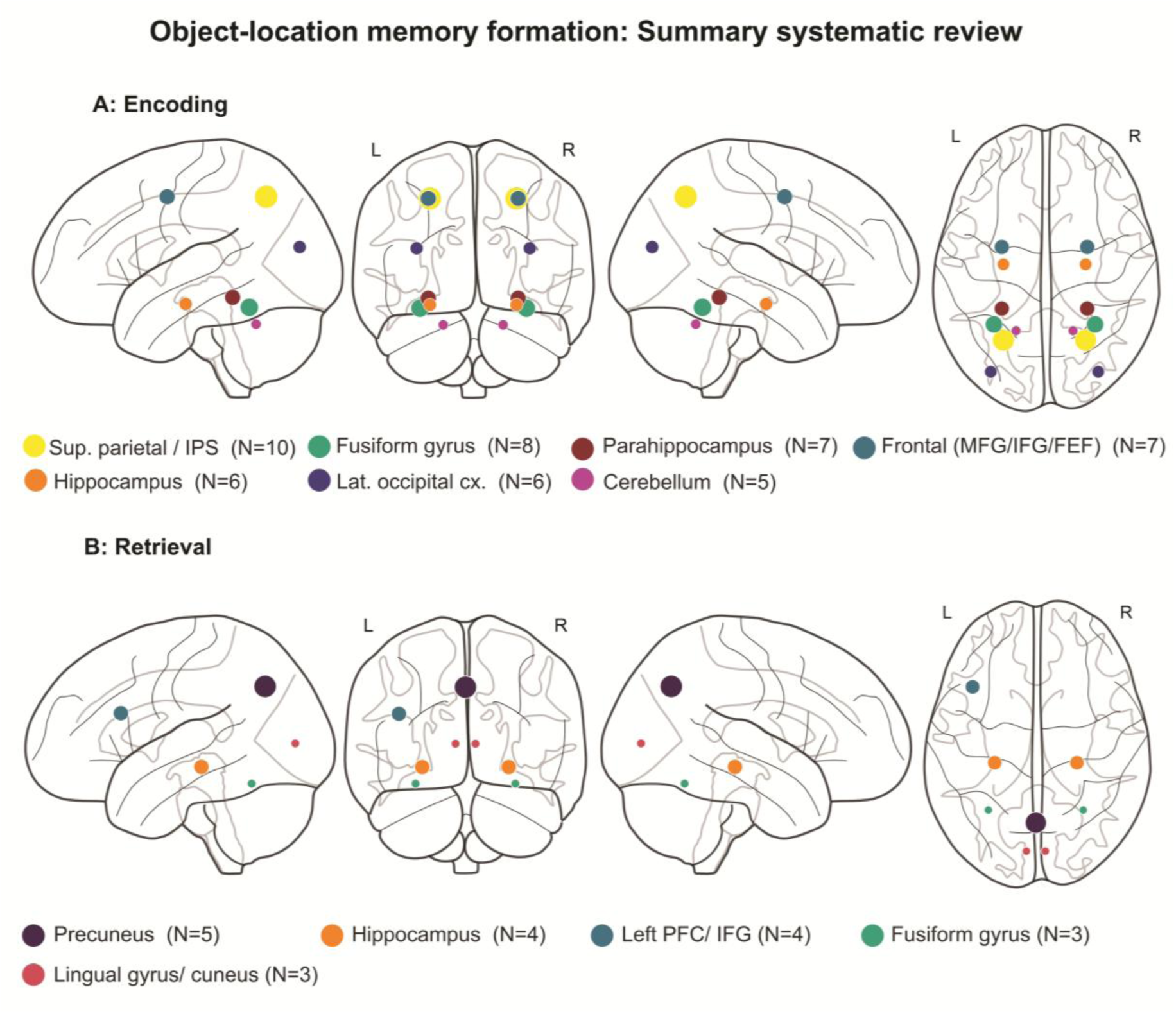
Schematic summary of systematic review results. Panel A. Glass brain visualization summarizing brain regions reported during object-location memory (OLM) encoding across the systematic review. Node size and color indicate the number of studies reporting activation in each region (N = 5a-10). Coordinates are displayed in MNI space. IPS = intraparietal sulcus; FEF = frontal eye fields; MFG = middle frontal gyrus; cx. = cortex. Panel B. Glass brain visualization summarizing brain regions reported during object-location memory (OLM) retrieval across the systematic review. Node size and color indicate the number of studies reporting activation in each region (N = 3-5). Coordinates are displayed in MNI space. OLM = object-location memory; PFC = prefrontal cortex; IFG = inferior frontal gyrus.

#### 3.2.2. Brain regions associated with OLM retrieval

A formal ALE for retrieval was not performed due to a lack of retrieval studies reporting coordinate data suitable for coordinate-based ALE meta-analysis. Although several included studies examined OLM retrieval, they generally did not report the whole-brain, task-versus-baseline peak coordinates that ALE requires: for example, activations were given by Brodmann-area label without coordinates (de Rover et al., 2008) or presented only as figures with no peak coordinates (Meulenbroek et al., 2010). The few usable retrieval foci (4 studies; Cansino et al., 2002, Slotnick, 2003, Ross & Slotnick, 2008, Kukolja et al., 2009) were too sparse to meet the minimum number of experiments required for an adequately powered ALE (Eickhoff et al., 2016; Müller et al., 2018). The retrieval findings are therefore reported through the qualitative synthesis instead.

##### Hippocampus

In contrast to encoding, hippocampal activation at retrieval was the most consistent finding across studies. For example, right hippocampal activation for correct spatial source retrieval was reported (Cansino, 2002) and replicated bilaterally with the left hippocampus active across age groups (Kukolja et al., 2009). This finding was confirmed via both whole-brain and ROI analysis (Ross & Slotnick, 2008). Right hippocampal activation during source retrieval was additionally identified in a multivariate analysis of an older adult cohort (Rabipour et al., 2020). Effective connectivity analysis further identified the right anterior hippocampus as the primary driver of retrieval connectivity in healthy older adults, with this pattern absent in MCI (Hampstead et al., 2016). An exception was Brodt et al. (2018), who found no significant hippocampal retrieval activation, interpreting this as evidence for rapid transfer of object-location representations to neocortical storage within 90 minutes of learning.

##### Precuneus

The precuneus was consistently activated during OLM retrieval. Bilateral precuneus activation increased progressively across recall repetitions, persisted beyond a 12-hour interval, and correlated with memory performance (Brodt et al., 2018). Precuneus activation at retrieval was additionally reported in Kukolja et al. (2009), Ross and Slotnick (2008), and Frings et al. (2006), the latter identifying the precuneus as the only region active during both encoding and retrieval in a conjunction analysis. The precuneus was further identified as the primary retrieval region in older adult cohorts (Rabipour et al., 2020, 2021).

##### Occipitotemporal cortex and prefrontal cortex

Several studies reported ventral-stream reactivation during retrieval. Fusiform and lingual/cuneus activation was greater for spatial than temporal associative retrieval (De Rover et al., 2008) and fusiform activation was identified during correct source retrieval (Kukolja et al., 2009; Ross & Slotnick, 2008). Occipitotemporal reactivation at retrieval, including lingual gyrus and occipital regions, was further supported by multivariate analyses (Rabipour et al., 2020, 2021).

Parahippocampal activation was also reported at retrieval (Cansino, 2002; De Rover et al., 2008). Additionally, left-lateralized prefrontal activation was associated with spatial source retrieval across multiple studies (Cansino, 2002; Ross & Slotnick, 2008; Slotnick et al., 2003). Prefrontal retrieval activation was also reported in multivariate analyses of older adult cohorts (Rabipour et al., 2020, 2021).

A schematic summary of the OLM retrieval is displayed in Figure 2B.

#### 3.2.3. Age-related differences (encoding and retrieval)

Older adults showed reduced activation in ventral and dorsal visual processing streams during encoding. A fusiform subsequent memory effect was found in young but not older adults (Kukolja et al., 2009), and reduced bilateral posterior fusiform, parahippocampal, and superior parietal activation was reported in older adults (Meulenbroek et al., 2010). In contrast, Hampstead et al. (2011) found that healthy older adults recruited the full encoding network, including bilateral fusiform, parahippocampal cortices, and hippocampus.

Beyond these activation differences, the pattern of hippocampal engagement also differed qualitatively between age groups. Young adults showed greater hippocampal activation for correct than incorrect source retrieval, while older adults showed the reverse pattern in the anterior subfield (Kukolja et al., 2009). At the same time, parahippocampal activation during encoding correlated more strongly with subsequent retrieval success in older adults than young adults (Kukolja et al., 2009).

Age-related differences in deactivation patterns were also observed. Most age-contrasts reflected reduced suppression of default mode network (DMN) regions during task performance, including medial PFC, ACC, and posterior cingulate (Kukolja et al., 2009; Meulenbroek et al., 2010). Alongside this, right inferior frontal gyrus showed above-baseline activation exclusively in older adults during retrieval (Meulenbroek et al., 2010). In clinical populations, mild cognitive impairment (MCI) patients showed reduced engagement of a left-lateralized frontoparietal encoding network and increased activation in frontal eye fields, associated with basic visual search (Hampstead et al., 2016). In the same paradigm, anterior hippocampal activation was selectively absent in MCI while posterior hippocampal activation was preserved (Hampstead et al., 2011). APOE4 carriers showed altered brain-behavior correlations, with performance linked to thalamus, supramarginal gyrus, and insula rather than the canonical MTL network (Rabipour et al., 2020). Additionally, sex-specific patterns of functional dedifferentiation were observed in older adults at risk for AD, with women showing less differentiated activation across source hit and source failure conditions than men (Rabipour et al., 2021).

### 3.3 ALE meta-analysis: OLM encoding

The primary ALE analysis of OLM encoding included 12 experiments from 12 studies, comprising 309 foci (mean 25.8 ± 33.4 foci per study), and reported from 234 participants (mean 19.5 ± 8.74 participants per study). Included studies, contrasts, and peak coordinates for each study are reported in supplementary table Table S1. Inference followed a cluster-level family-wise error correction procedure (p-FWE < 0.05; cluster-forming threshold p < 0.001 uncorrected; 10,000 Monte Carlo permutations), yielding a minimum cluster-size threshold of 856 mm³. Per-experiment ALE spatial uncertainty kernels (FWHM) ranged from 8.85 to 9.76 mm (median 9.50 mm).

Two clusters survived cluster-level FWE correction (total suprathreshold volume = 3,648 mm³). Cluster locations, volumes, peak ALE values, and anatomical labels were obtained from the Talairach Daemon (default in GingerALE) and cross-referenced against the Harvard-Oxford and Jülich-Brain atlases for cerebral peaks and the SUIT atlas for cerebellar peaks; these are reported in Table 3 and visualized in Figure 3.

**Figure 3.**
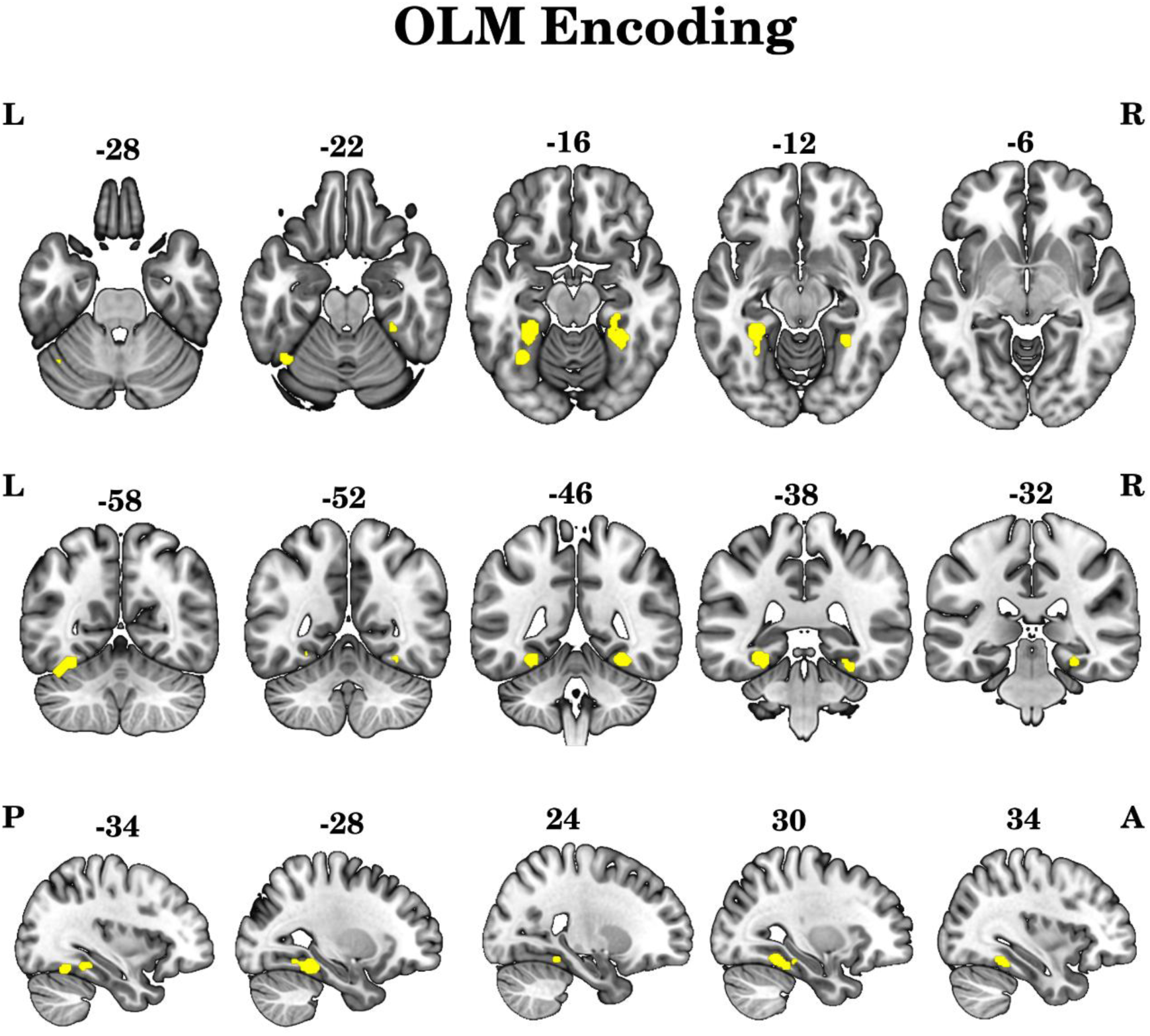
Convergent activation during OLM encoding. Thresholded ALE map (cluster-level FWE-corrected p < 0.05; cluster-forming threshold p < 0.001 uncorrected) overlaid on the MNI152 template and visualized using MRIcroGL. Rows display axial, coronal, and sagittal views, respectively. Two bilateral clusters are significant: a left-sided cluster encompassing the fusiform gyrus, parahippocampal gyrus, inferior temporal gyrus, and cerebellum, and a right-sided cluster involving the fusiform gyrus, parahippocampal gyrus, and cerebellum.

**Table 3.**
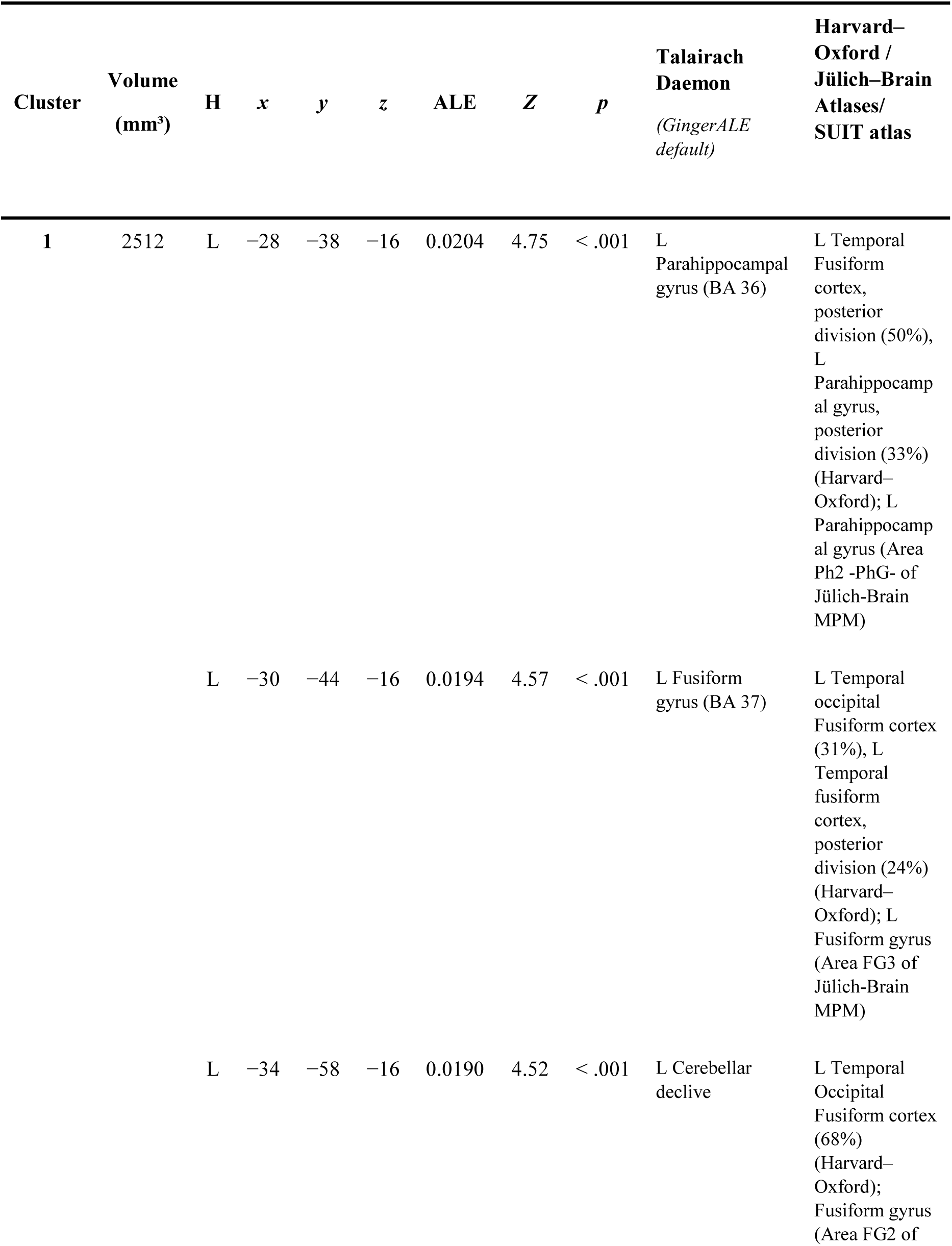
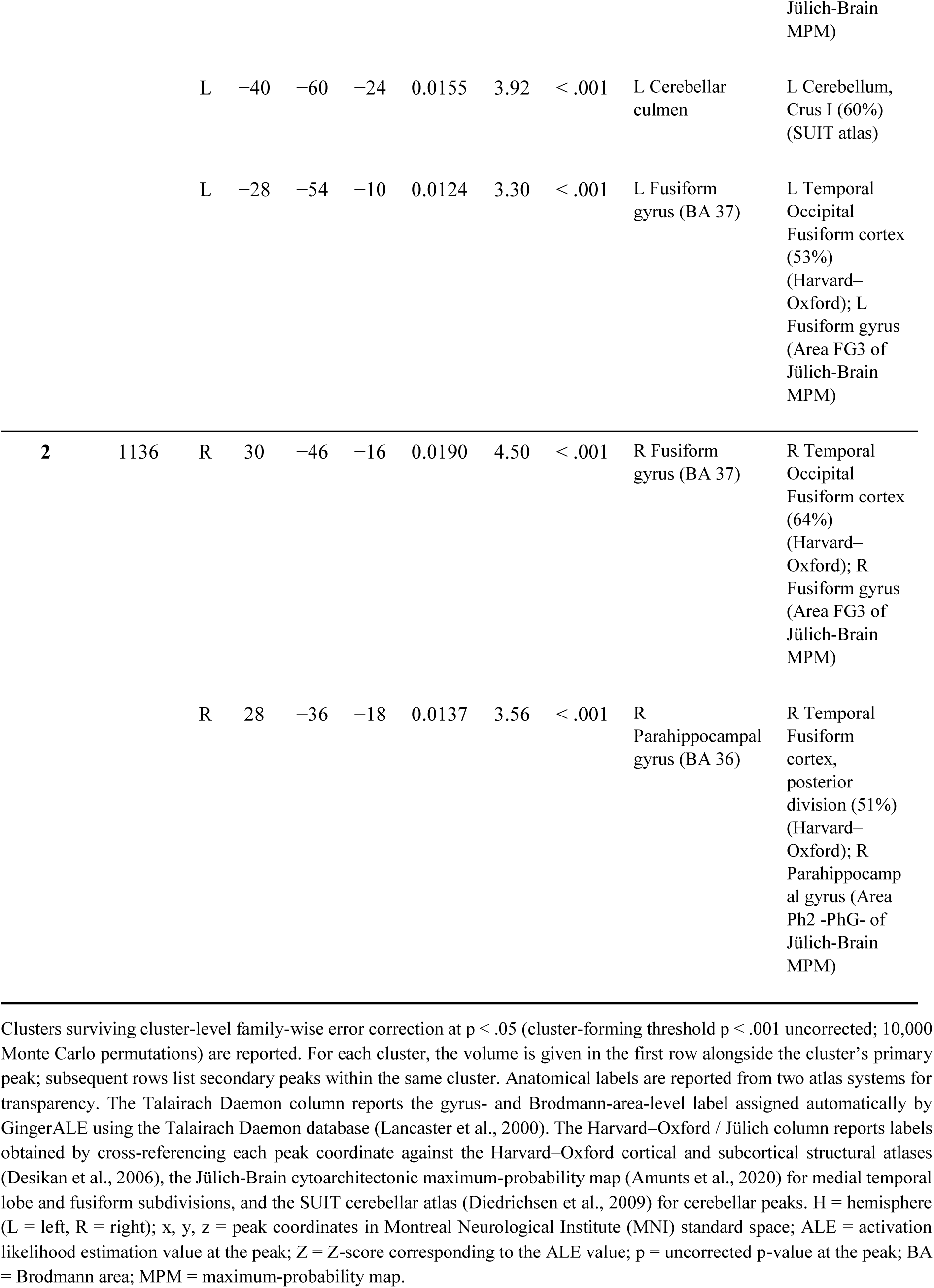
Clusters of convergent activation identified by the activation likelihood estimation meta-analysis of object–location memory encoding.

Cluster 1 was located in the left occipitotemporal-cerebellar region, with its peak at MNI (-28, -38, -16) (peak ALE = 0.02; Z = 4.75; p < 0.001; cluster volume = 2,512 mm³). The cluster spanned the left fusiform gyrus (Brodmann area 37), left parahippocampal gyrus (Brodmann area 36), and left cerebellum, with extension into the neighboring left inferior temporal gyrus (Brodmann area 20). Eight of the 12 included experiments contributed foci to this cluster: Abdelmotaleb (2025; 3 foci), Hampstead (2011; 4 foci), Gillis (2016; 2 foci), Sommer et al. (2005, LM; 1 focus), Sommer et al. (2005, Neuropsychologia; 1 focus), Cansino (2002; 1 focus), Gould (2003; 1 focus), and Ross and Slotnick (2008; 1 focus).

Cluster 2 was located in the right occipitotemporal-cerebellar region, with its peak at MNI (30, -46, -16) (peak ALE = 0.019; Z = 4.50; p < 0.001; cluster volume = 1,136 mm³). The cluster spanned the right parahippocampal gyrus and right fusiform gyrus, with anatomical labeling indicating contributions from Brodmann areas 37, 36, 35, and 20. Four of the 12 included experiments contributed foci to this cluster: Abdelmotaleb (2025; 2 foci), Hampstead (2011; 2 foci), Gillis (2016; 1 focus), and Gould (2003; 1 focus).

The convergent activation pattern was therefore broadly bilateral, with left- and right-hemisphere clusters showing comparable peak coordinates along the y- and z-axes. The two clusters were supported by 8 of 12 (67%) and 4 of 12 (33%) contributing experiments, respectively. Subpeak coordinates and Z-scores for both clusters are reported in Table 3.

A formal coordinate-based ALE comparison of age groups was not performed because the number of age-comparison experiments fell below the threshold for adequate ALE power, and the available contrasts pooled across encoding and retrieval phases (see Discussion).

## 4. Discussion

In this systematic review and coordinate-based meta-analysis, we investigated task-dependent fMRI activation during OLM formation in healthy adults, with a primary focus on the encoding of object-location associations and a complementary analysis of OLM retrieval. A secondary objective was to explore activation differences between younger (18-45 years) and older (55-80 years) adults. To our knowledge, this is the first systematic review and coordinate-based meta-analysis dedicated specifically to OLM. Overall, 21 studies with 637 participants were included in the systematic review, and 12 studies met the criteria for a coordinate-based meta-analysis, contributing 309 foci from 234 participants to an ALE of OLM encoding activity. In the systematic review, we showed that OLM encoding consistently recruited the bilateral fusiform gyrus and parahippocampal cortex, with additional but less convergent engagement of parietal and prefrontal regions across individual studies, whereas OLM retrieval was characterized by activation of the hippocampus and precuneus. The ALE meta-analysis for encoding of OLM yielded two bilateral occipitotemporal clusters that spanned the fusiform and parahippocampal gyri and extended to the occipitotemporal-cerebellar border. Age-related differences were sparsely reported, but pointed toward (a) a pattern resembling the posterior-anterior shift in aging (PASA; Davis et al., 2008), with reduced activation in posterior fusiform, parahippocampal, and superior parietal regions alongside increased activation in prefrontal and midline regions, and (b) reversed hippocampal pattern at retrieval, with older adults showing greater activation for retrieval failures than successes. Taken together, OLM encoding appears to be processed primarily within the ventral visual-to-medial-temporal processing stream, whereas retrieval additionally recruits the hippocampus and precuneus. Age-related differences point not to uniform hypoactivation but to distinct alterations along the same pathways.

### 4.1. OLM Encoding

The fusiform gyrus and parahippocampal cortex were among the most consistently reported encoding regions in the systematic review and this pattern was replicated by the ALE, which confirmed two bilateral clusters. The clusters fell within the posterior temporo-occipital fusiform gyrus, a key component of the object-selective ventral visual stream (Kanwisher et al., 1997; Weiner & Zilles, 2016), and in the parahippocampal cortex, which lies at the interface between high-level visual cortex and hippocampal memory systems and is implicated in contextual associative processing (Aminoff et al., 2013). Across studies, fusiform activation was the dominant ventral stream response. It showed good test-retest reliability (Abdelmotaleb et al., 2025), a robust subsequent-memory effect driven by perceptual engagement at encoding (Cansino, 2002), and a parametrically graded relationship with memory confidence (Sommer, Rose, Gläscher, et al., 2005; Sommer, Rose, Weiller, et al., 2005). Brodt et al. (2018) further reported significant learning-induced microstructural changes in the fusiform gyri during object-location learning relative to a control condition. Parahippocampal cortex was engaged during successful spatial-source encoding (Ross & Slotnick, 2008), and during the associative binding of object-location associations (Gillis et al., 2016; Hampstead et al., 2011). Its activation also predicted successful contextual memory performance (Kukolja et al., 2009) and showed a partially dissociable and anatomically specific profile: posterior parahippocampal cortex predicted memory for both the object and location components, whereas more anterior medial-temporal clusters were specifically predictive of location recall (Sommer, Rose, Gläscher, et al., 2005), and left parahippocampal cortex was engaged more strongly during encoding than during retrieval (Gould et al., 2003).

In the only previous systematic review on OLM, Zimmermann & Eschen (2017) concluded from PET and fMRI studies that the inferior temporal/fusiform gyrus is specialized for object processing in small space episodic OLM. Here, we provide the first quantitative whole-brain confirmation of this conclusion. Additionally, the ALE convergence aligns closely with prior meta-analytic evidence. Both Cona and Scarpazza (2019) and Torres-Morales and Cansino (2024) reported parahippocampal and ventral occipitotemporal peaks in closely overlapping locations, supporting the reproducibility of these regions across independent analyses.

Additionally, this ventral-dominant encoding involvement is consistent with the influential three-component model of Postma and colleagues (2004, 2008), in which successful OLM requires object identity processing via the ventral visual stream, spatial-location processing via the dorsal visual stream, and binding within the hippocampus and broader MTL. The meta-analysis results support the ventral object-identity component as the most spatially consistent element of OLM encoding. Although the model also predicts dorsal stream involvement, parietal and frontal activation did not survive ALE thresholding despite being reported across multiple individual studies (e.g., Abdelmotaleb et al., 2025; Gillis et al., 2016; Gould et al., 2003; Hampstead et al., 2011; Sommer, Rose, Gläscher et al., 2005; Sommer, Rose, Weiller et al., 2005). Rolls (2024) proposed that spatial location information reaches the hippocampus primarily through the ventromedial occipitotemporal pathway involving ventromedial visual regions (“where”) and the parahippocampus (“what”), rather than the posterior parietal cortex. Consequently, the dorsal parietal stream might contribute to self-motion updating and coordinate transformations rather than primary spatial input (Rolls, 2024; Wolbers et al., 2008). The present results converge on precisely these ventromedial regions without parietal convergence, providing additional support for a ventral-dominant input pathway during OLM encoding.

Nevertheless, connectivity evidence suggests that both streams interact during OLM formation: effective connectivity from dorsal parietal to ventral fusiform cortex increased over the course of object-location learning (Büchel et al., 1999). Additionally, activity in parietal regions including the IPS and precuneus and frontal regions including the MFG has been identified across multiple individual studies (Abdelmotaleb et al., 2025; Frings et al. 2006; Gillis et al., 2016; Gould et al., 2003; Hampstead et al., 2011). These frontal and parietal activations may reflect general spatial attention and working memory processes that are not directly involved in hippocampal binding (Cona & Scarpazza, 2019; Gillis et al., 2016; Hales & Brewer, 2013). Consistent with this interpretation, Kim and Voss (2019) found that less coupling of frontoparietal regions during encoding predicted better OLM performance, suggesting that excessive frontoparietal engagement may not be beneficial for OLM encoding.

Both ALE clusters nominally extended into the cerebellum, but this should be interpreted with caution. Cross-atlas comparison reassigned most peaks labeled cerebellar by the Talairach Daemon to ventral occipitotemporal cortex under the Harvard-Oxford and Jülich atlases, with only one peak (−40, −60, −24, Crus I) falling within cerebellar tissue. Where primary studies reported cerebellar activation (5 of 12), they attributed it to motor or domain-general processes rather than object-location binding (Frings et al., 2006; Zimmermann & Eschen, 2017). We therefore do not regard the cerebellar extension as a core component of the OLM encoding network, although a contributory role cannot be excluded. Object-to-location binding is thought to be supported by MTL regions, with the hippocampus as the principal contributor (Mayes et al., 2007; Postma et al., 2008). Hippocampal activity was reported in individual studies (Abdelmotaleb et al., 2025; Gillis et al., 2016; Gould et al., 2003; Hampstead et al., 2011), but did not survive ALE thresholding at encoding, in line with prior coordinate-based meta-analyses (Cona & Scarpazza, 2019; Torres-Morales & Cansino, 2024). In contrast, lesion-based literature synthesized by Postma et al. (2008) and Zimmermann & Eschen (2017), drawing heavily on patient data, has emphasized the contribution of the hippocampus in their reviews. This discrepancy most likely reflects the well-recognized lower sensitivity of fMRI relative to lesion studies to medial-temporal involvement (Aggleton & Brown, 2006; Zimmermann & Eschen, 2017), rather than a genuine absence of hippocampal contribution. Two design features of the present review are also relevant. First, the review focused specifically on OLM and excluded broader spatial memory and navigation paradigms, which may engage the hippocampus differently (Sagi et al., 2012). Second, the separation of encoding and retrieval means that hippocampal effects primarily expressed during retrieval would not be captured in the encoding analysis. This is supported by the finding that the hippocampus was among the most consistent retrieval regions in the systematic review (see below).

### 4.2. OLM Retrieval

Because too few studies reported usable retrieval coordinates, an ALE meta-analysis was not feasible, and the retrieval findings are based on the systematic review. The hippocampus emerged as the most consistent retrieval region. Right hippocampal activation was reported for correct spatial-source retrieval (Cansino, 2002), and left hippocampal engagement was observed across age groups during successful contextual retrieval (Kukolja et al., 2009). The left hippocampus also showed greater involvement in spatial-source retrieval in both whole-brain and ROI-based analyses (Ross & Slotnick, 2008). The principal exception was Brodt and colleagues (2018), who reported no significant hippocampal activation at either encoding or retrieval of object-location associations in their whole-brain analysis, but identified right hippocampal activation during encoding in a region-of-interest analysis. Hippocampal retrieval engagement is consistent with the reinstatement hypothesis, in which successful retrieval re-engages encoding regions (for a review, see Danker & Anderson, 2010; Rugg et al., 2008). This interpretation was supported by the effective-connectivity study of object-location associations by Hampstead and colleagues (2016), in which the hippocampus was the main driver during object-location retrieval.

The precuneus was likewise consistently engaged during retrieval (Kukolja et al., 2009; Rabipour et al., 2020, 2021), and was the only region active during both encoding and retrieval in a conjunction analysis (Frings et al., 2006). Bilateral precuneus activation increased across recall repetitions, persisted beyond a 12-hour interval, and correlated with memory performance (Brodt et al., 2018). It was also identified as the primary retrieval region in older adults (Rabipour et al., 2020). As a node of the posterior-medial network supporting contextual reconstruction, the precuneus appears to support the internal visualization and spatial reinstatement of the remembered layout (Cavanna & Trimble, 2006; Ranganath & Ritchey, 2012). Its rapid and sustained engagement further suggests that it may serve as an early neocortical store for object-location representations, complementing the hippocampus (Schott et al., 2018).

Two additional systems supported retrieval. First, ventral occipitotemporal regions were reactivated, with greater fusiform and lingual activity during spatial than temporal associative retrieval (De Rover et al., 2008) and fusiform and middle/inferior temporal cortices during correct source retrieval (Kukolja et al., 2009; Ross & Slotnick, 2008; Cansino et al., 2002). Because these were the same ventral regions that dominated encoding, their reactivation is consistent with the reinstatement of encoded perceptual and object information. Second, the left lateral prefrontal cortex was consistently associated with spatial-source retrieval across studies (Cansino, 2002; Ross & Slotnick, 2008; Slotnick et al., 2003). This region is thought to support controlled retrieval processes, including cue specification, source monitoring, and response selection (Badre & Wagner, 2007). Its left lateralization is consistent with a contribution of verbal and semantic recoding to source memory.

### 4.3. Age-related differences

The systematic review identified a pattern of both hypo- and hyperactivation in older compared with young adults during OLM processing. Older adults showed reduced activation in posterior cortical regions, including fusiform gyrus, parahippocampal cortex, and superior parietal cortex (Kukolja et al., 2009; Meulenbroek et al., 2010). In contrast, increased activation was observed in the right inferior frontal gyrus, which showed above-baseline activation exclusively in older adults during retrieval (Meulenbroek et al., 2010), and in the midline regions including medial prefrontal cortex, anterior cingulate and posterior cingulate cortex (Kukolja et al., 2009; Meulenbroek et al., 2010). However, these findings are based on a small number of studies (3 of 21 permitted a within-study age comparison, supplemented by four studies that examined older adults or clinical samples without a young adult control group) with direct age-group comparison and should be interpreted with caution.

This pattern of posterior reductions coupled with anterior increases might resemble the PASA (Davis et al., 2008), in which age-related reductions in occipitotemporal activity are accompanied by increased frontal recruitment. Whether such frontal increases reflect functional compensation or neural dedifferentiation remains debated (Cabeza, 2002; Cabeza et al., 2018), and in the present review the prefrontal hyperactivation rests on a single study (Meulenbroek et al., 2010), limiting conclusions about its functional significance. At the structural level, aging is associated with atrophy of the hippocampus and MTL, including the parahippocampal and fusiform gyri, which correlates with connectivity loss and impaired memory performance across the lifespan (Blinkouskaya et al., 2021; Fjell & Walhovd, 2010; Snytte et al., 2024; Squire et al., 2004). The present review extends this picture by suggesting specific functional alterations in OLM encoding and retrieval. Older adults showed reduced activation in ventral and dorsal processing streams during encoding, with a fusiform subsequent memory effect present in younger but not older adults (Kukolja et al., 2009), and reduced bilateral fusiform, parahippocampal, and superior parietal activation (Meulenbroek et al., 2010). These findings suggest that age-related OLM impairment may arise upstream of the MTL, in perceptual processing related to the ventral encoding pathway identified in the ALE. Additionally, older adults showed increased activation in midline regions typically suppressed during task performance, including medial prefrontal cortex, anterior cingulate, and posterior cingulate cortex (Kukolja et al., 2009; Meulenbroek et al., 2010). This reduced suppression is a well-documented finding in the cognitive aging literature and has been associated with poorer memory performance in older adults (Park & Reuter-Lorenz, 2009; Stern et al., 2019). Beyond these activation differences, the pattern of hippocampal engagement also differed between age groups. Young adults showed greater hippocampal activation for correct than incorrect source retrieval, while older adults showed the reverse pattern in the anterior subfield (Kukolja et al., 2009). At the same time, parahippocampal activation during encoding correlated more strongly with subsequent retrieval success in older than younger adults (Kukolja et al., 2009).

Age-related differences in OLM-related activation involved both quantitative reductions in posterior encoding regions and a qualitative reorganization of hippocampal engagement at retrieval, though these findings rest on a small and heterogeneous set of studies and should be treated as preliminary.

Notably, these age-related reductions were not universal. Hampstead et al. (2011) found that healthy older adults recruited the full encoding network, including bilateral fusiform and parahippocampal cortex and the hippocampus, suggesting that the posterior cortical reductions reported elsewhere may depend on task demands, sample characteristics, or individual differences in brain maintenance rather than reflecting a uniform consequence of aging (Cabeza et al., 2018). In clinical populations, the pattern was more pronounced (Hampstead et al., 2011, 2016). For example, MCI patients showed reduced engagement of a left lateralized fronto-parietal encoding network and increased activation in regions linked to basic visual search, suggesting reduced goal-directed encoding (Hampstead et al., 2016). APOE4 carriers showed altered brain-behavior relationships, with performance linked to thalamus, supramarginal gyrus, and insula rather than the MTL network (Rabipour et al., 2020). OLM impairments have been reported among the earliest cognitive signs of AD, emerging at preclinical stages and positioning OLM as a candidate behavioral marker of early hippocampal dysfunction (De Sousa et al., 2020; Hampstead et al., 2018). The present findings provide a neural framework for this. If OLM encoding relies on ventral visual and parahippocampal-to-hippocampal processing, then early pathology in the hippocampus and entorhinal cortex may selectively disrupt OLM. OLM-based fMRI paradigms could therefore be explored as markers of early AD-related network dysfunction.

### 4.4 Limitations

Several limitations should be considered when interpreting our findings: First, the methodological heterogeneity across included studies. Paradigms, contrast types, correction methods, and control conditions varied widely across the included studies, from uncorrected voxelwise thresholds (Cansino, 2002; Kukolja et al., 2009; Sommer, Rose, Weiller et al., 2005) to cluster- and voxel-level family-wise-error correction, and from low-level fixation baselines to demanding active control tasks. This heterogeneity limits the comparability of individual findings and may have reduced the likelihood of ALE convergence for regions sensitive to paradigm type. However, by restricting inclusion to OLM-specific paradigms and separating encoding from retrieval, this review directly addresses the heterogeneity in how OLM has been operationalized and assessed.

Second, the small sample sizes in primary studies. Eleven of the 21 included studies analyzed 15 or fewer participants, and the median sample was approximately 20, which inflates the risk of unreliable peak localization and reduces sensitivity to modest MTL effects. However, the ALE algorithm down-weights smaller studies, and the main ventral-stream finding was independently reported by two previous meta-analyses (Cona & Scarpazza, 2019; Torres-Morales & Cansino, 2024), supporting its robustness.

Third, the limited number of experiments for meta-analytic synthesis. The encoding ALE rested on 12 experiments, below the approximately 17–20 generally recommended for adequately powered ALE (Eickhoff et al., 2016; Müller et al., 2018). This recommendation depends on expected effect size, and results from smaller analyses should be interpreted with caution. Additionally, a number of informative studies could not contribute coordinates because they employed connectivity, multivariate, or parcellation-based approaches (Büchel et al., 1999; Kim & Voss, 2019; Rabipour et al., 2020, 2021; Rolls et al., 2024), though these were retained in the qualitative synthesis. Too few studies were available to perform either a retrieval ALE or a coordinate-based age comparison, leaving those questions to the systematic review.

### 4.5 Implications and future directions

Several methodological priorities for future studies follow from the limitations of the currently available evidence. First, studies should report standard whole-brain encoding > control contrasts alongside subsequent-memory analyses, so that meta-analytic pooling is not confounded by contrast type. Second, adequately powered age-group designs with matched OLM tasks and explicitly reported encoding-phase age contrasts would address a gap left by prior reviews. Third, clearer specification of whether a paradigm targets allocentric object-in-location binding or coarse spatial source would help resolve a distinction that appears to recruit partly distinct networks (Zimmermann & Eschen, 2017).

The present findings also carry translational implications. By identifying the ventral occipitotemporal-medial pathway as the most spatially consistent substrate of OLM formation, the results may inform targeted network-level interventions. NIBS techniques such as transcranial direct current stimulation, temporal interference, or transcranial magnetic stimulation could be used to directly target specific nodes of the OLM encoding network to modulate performance in both healthy aging and early neurodegenerative diseases (Bjekić et al., 2019; Flöel et al., 2012). Future studies combining task fMRI with effective- or functional-connectivity analyses are needed to pinpoint the specific connectivity changes that support successful OLM encoding, within the ventral occipitotemporal–MTL pathway, and between this pathway and parietal/prefrontal regions. This direction is particularly relevant because OLM impairments are among the earliest cognitive signs of AD and have been proposed as a behavioral marker of early hippocampal dysfunction (De Sousa et al., 2020; Hampstead et al., 2018).

## 5. Conclusions

This first systematic review and coordinate-based meta-analysis of OLM identifies the ventral visual-to-medial-temporal pathway as the most spatially consistent substrate of OLM encoding and reveals an encoding-retrieval dissociation, with the hippocampus and precuneus emerging primarily at retrieval. Current OLM models do not account for these phase-dependent differences or the age-related alterations observed along this pathway. A clearer characterization of the neural substrates of OLM formation across the adult lifespan is likely to be of value for both basic and clinical research. The ventral encoding pathway identified here may inform both the theoretical understanding of OLM and future translational approaches, including targeted network-level interventions such as noninvasive brain stimulation to counteract cognitive decline in aging and neurodegenerative disease.

## Data and code availability

The data supporting this study’s findings are available from the corresponding authors upon reasonable request.

## Author contributions

Anna Elisabeth Fromm: literature research, data extraction, analysis, figure preparation, writing first draft, writing (review and editing)

Mohamed Abdelmotaleb: literature research, data extraction, analysis, figure preparation, writing first draft, writing (review and editing)

Freya Olschewski: literature research, data extraction, writing (review and editing)

Jakub Limanowski: writing - review & editing, supervision, project administration

Marcus Meinzer: writing - review & editing, supervision, project administration, funding acquisition

Agnes Flöel: conceptualization, writing - review & editing, supervision, project administration, funding acquisition

Daria Antonenko: conceptualization, methodology, investigation, writing – original draft, writing - review & editing, supervision, project administration, funding acquisition

All authors approved the paper.

## Supplementary Material

**Table S1.** Activation foci reported by the 12 functional magnetic resonance imaging studies included in the encoding ALE meta-analysis of object-location memory.

| # | Study | Contrast | N | Foci (n) | Peak coordinates (MNI x, y, z) |
| --- | --- | --- | --- | --- | --- |
| 1 | Abdelmotaleb (2025) | Learning > Control | 20 | 34 | (30, -46, -14); (30, -36, -22); (36, -14, -22); (-32, -58, -16); (-26, -36, -18); (-30, -48, -16); (-34, -86, 26); (-40, -86, 18); (-26, -72, 30); (32, -78, 4); (40, -74, 22); (40, -84, 14); (12, 20, -4); (6, 12, -8); (20, 10, -8); (34, -26, 68); (16, -24, 76); (44, -22, 50); (-8, 14, -4); (16, 30, -14); (26, 32, -14); (8, 38, -14); (-6, -56, 50); (8, -54, 46); (-10, -70, 54); (-16, -10, 56); (-28, -14, 54); (-20, 0, 56); (-28, 26, -4); (-32, 18, -2); (-32, 24, -14); (-40, -60, -4); (-48, -64, -6); (-46, -64, 2) |
| 2 | Brodt (2018) | Encoding correct > incorrect (subsequent memory, Table S9) | 39 | 5 | (-38, 6, -36); (-44, 24, -24); (26, -78, -36); (-10, 62, 24); (36, -24, -10) |
| 3 | Frings (2006) | Encoding vs. perceptual/motor control (ENC1 > COMP1) | 13 | 2 | (-24, -69, 57); (15, -66, 54) |
| 4 | Kukolja (2009) | CorSCE > FalSCE (main effect, encoding) | 35 | 1 | (-42, -48, -12) |
| 5 | Sommer, Rose, Gläscher et al., (2005) | Encoding - object-cue subsequent memory | 15 | 18 | (3, -93, 21); (39, -72, 24); (33, -63, 48); (-9, -81, 48); (-30, -57, 45); (21, -63, -9); (-18, -60, -6); (45, -45, -18); (45, -48, -18); (-42, -63, -24); (-42, -48, -27); (21, -63, -9); (-36, -33, -15); (-27, 3, 48); (30, 3, 51); (-45, 27, 15); (-24, -9, -27); (30, -6, -27) |
| 6 | Sommer, Rose, Weiller et al., (2005) | Encoding - parametric subsequent memory | 15 | 16 | (3, -96, 12); (42, -81, 21); (27, -60, 54); (-15, -78, 54); (-33, -60, 36); (39, -39, -24); (42, -57, -18); (-45, -54, -21); (27, -57, -9); (-33, -42, -12); (-27, -6, 45); (36, -3, 48); (-42, 54, 0); (-51, 21, -3); (51, 36, 27); (-6, 24, 36) |
| 7 | Cansino (2002) | Correct > Incorrect subsequent source memory (encoding) | 17 | 16 | (-20, 51, 44); (-3, 40, 61); (-40, 7, 16); (58, 1, 55); (-50, 3, 57); (-39, -3, 64); (2, -30, 68); (-46, -40, 36); (-61, -56, 6); (-44, -54, 4); (55, -73, 2); (31, -78, -9); (42, -49, -14); (-40, -59, -25); (-14, 17, -19); (-10, -60, -19) |
| 8 | Gillis (2016) | Object+Location novel > Object+Location repeated | 15 | 96 | (-29, 66, 3); (45, 20, 26); (43, 32, 18); (-40, 24, 27); (50, 5, 31); (-44, 4, 34); (45, 28, 23); (51, 43, 15); (-46, 23, 30); (-47, 9, 40); (-36, 51, 14); (2, 19, 47); (-3, 19, 46); (-14, 4, 61); (32, -1, 58); (-27, -3, 58); (37, 0, 52); (-35, -2, 59); (-52, 6, 38); (34, 23, -3); (-29, 21, -1); (10, 29, 26); (-6, 30, 32); (12, 12, 2); (-8, 15, 3); (31, 4, -4); (-21, 7, -3); (11, -17, 7); (9, -15, 7); (-8, -15, 8); (-11, -17, 5); (19, -28, 5); (-15, -23, 6); (9, -32, -6); (-5, -28, -5); (30, -29, -15); (-24, -20, -13); (-30, -25, -14); (21, -23, -24); (-22, -24, -26); (14, -9, -22); (-15, -9, -26); (20, -11, -25); (-19, -10, -27); (28, -41, -18); (-26, -42, -18); (38, -8, -28); (27, -38, -15); (35, -67, -14); (-34, -7, -32); (-26, -38, -16); (-38, -67, -13); (50, -58, -20); (-52, -56, -16); (15, -44, -6); (-14, -45, -8); (-11, -33, 49); (6, -34, 24); (-5, -36, 25); (9, -50, 8); (-6, -49, 9); (-60, -12, 19); (-59, -20, 50); (17, -69, 58); (-30, -36, 68); (-10, -71, 54); (37, -46, 43); (31, -61, 46); (24, -70, 41); (-43, -37, 38); (-36, -55, 46); (-25, -68, 42); (-46, -41, 36); (33, -75, 44); (-39, -68, 48); (8, -68, 51); (-7, -72, 47); (17, -97, 16); (-18, -96, 16); (60, -58, -8); (-53, -63, -3); (31, -86, 16); (-28, -86, 17); (26, -92, -10); (-15, -94, -10); (6, -82, -2); (-8, -92, -1); (35, -86, -10); (-38, -86, -12); (18, -70, -16); (-12, -71, -12); (19, -92, -9); (-11, -96, -10); (21, -58, -26); (-26, -58, -23); (1, -21, -27) |
| 9 | Gould (2003) | Zero-order encoding > rest (Table 2i) | 23 | 14 | (-29, 8, 47); (15, -58, 62); (-7, 23, 36); (-36, -60, -18); (32, -86, 2); (-46, 27, 15); (28, -73, 49); (51, -75, -1); (34, -74, 34); (-12, -12, 1); (-41, 1, 52); (32, 13, 45); (31, -50, -15); (-31, -79, 33) |
| 10 | Hales & Brewer (2013) | Correct Location > Incorrect Location (cue period) | 14 | 3 | (38, 40, 9); (-40, -63, -34); (4, 22, 38) |
| 11 | Hampstead (2011) | HEC Novel > Repeated (Supplementary Table 1) | 16 | 93 | (-4, 53, -16); (-36, 31, -11); (29, 33, -12); (13, 25, -17); (-4, 4, 52); (-4, 20, 41); (9, 17, 43); (-20, 7, 60); (-22, 5, 49); (25, 5, 48); (-45, 25, 31); (47, 29, 22); (-48, 34, 9); (-51, 19, 21); (52, 36, 8); (-41, 23, 23); (43, 26, 19); (-35, 18, 3); (37, 18, 2); (-45, 5, 23); (-25, -5, 49); (44, 7, 25); (29, -5, 49); (-49, -2, 48); (50, -7, 50); (-36, -34, 43); (-50, -25, 35); (42, -32, 42); (-4, -54, 15); (-4, -46, 48); (7, -54, 15); (7, -46, 48); (-8, -34, 29); (6, -33, 29); (-5, -49, 8); (8, -49, 8); (-12, -69, 56); (-28, -62, 54); (21, -66, 53); (28, -56, 50); (-37, -45, 41); (-26, -69, 33); (32, -45, 43); (26, -62, 45); (-29, -85, 30); (32, -80, 29); (-16, 7, 6); (18, 6, 5); (-7, -9, 2); (8, -10, 2); (-6, -16, 9); (8, -15, 9); (-15, -29, 3); (17, -29, 2); (-33, -9, -27); (27, -9, -31); (-21, -9, -26); (17, -8, -22); (-28, 1, -21); (-26, -6, -14); (19, -3, -19); (-22, -13, -20); (20, -11, -20); (25, -14, -19); (-30, -21, -14); (32, -21, -13); (-26, -32, -7); (-16, -36, 1); (30, -31, -7); (17, -35, 2); (-27, -27, -17); (-17, -32, -9); (23, -42, -14); (29, -21, -23); (-28, -53, -11); (-30, -39, -14); (26, -33, -16); (33, -58, -14); (-31, -43, -18); (-35, -54, -15); (29, -60, -13); (33, -43, -18); (-45, -58, -10); (48, -58, -11); (-33, -89, 13); (33, -79, 14); (-43, -80, -5); (40, -80, - |

| # | Study | Contrast | N | Foci<br>(n) | Peak coordinates (MNI x, y, z) |
| --- | --- | --- | --- | --- | --- |
|  |  |  |  |  | 7); (-32, -85, 30); (34, -83, 28); (-28, -73, -7);<br>(26, -73, -13); (0, -56, -27) |
| 12 | Ross & Slotnick<br>(2008) | enc-hit-hits > enc-hit-misses<br>(subsequent source memory,<br>Contrast A) | 12 | 11 | (-25, -43, -10); (62, -16, 47); (-53, -19, 54);<br>(46, -30, 56); (38, -38, 57); (-46, -32, 58); (36,<br>-52, 64); (-26, -57, -7); (-50, -68, 12); (-48, -<br>80, 4); (-44, -82, 9) |
| **Total** |  |  | **234** | **309** |  |

